# The distinct translational landscapes of Gram-positive and Gram-negative bacteria

**DOI:** 10.1101/2023.05.25.542305

**Authors:** Owain J. Bryant, Filip Lastovka, Jessica Powell, Betty Y-W Chung

**Affiliations:** Department of Pathology, University of Cambridge, Tennis Court Road, Cambridge, CB2 1QP, United Kingdom; Current address: Centre for Structural Biology, National Cancer Institute, 21702, Frederick, Maryland, USA

**Author notes:** Corresponding author *Betty Y-W Chung Department of Pathology, University of Cambridge, Tennis Court Road, Cambridge, CB2 1QP, United Kingdom. Contributed Equally (Owain J. Bryant, Filip Lastovka).

## Abstract

Translational control in pathogenic bacteria is fundamental to gene expression and affects virulence and other infection phenotypes. We used an enhanced ribosome profiling protocol coupled with parallel transcriptomics to capture accurately the global translatome of two evolutionarily distant pathogenic bacteria – the Gram-negative bacterium *Salmonella* and the Gram positive bacterium *Listeria* We find that the two bacteria use different mechanisms to translationally regulate protein synthesis. In *Salmonella,* in addition to the expected correlation between translational efficiency and *cis*-regulatory features such as Shine-Dalgarno (SD) strength and RNA secondary structure around the initiation codon, our data reveal an effect of the 2^nd^ and 3^rd^ codons, where the presence of tandem lysine codons (AAA-AAA) enhances translation in both *Salmonella* and *E. coli*. Strikingly, none of these features are seen in efficiently translated *Listeria* transcripts. Instead, approximately 15% of efficiently translated *Listeria* genes exhibit 70S footprints seven nt upstream of the authentic start codon, suggesting that these genes may be subject to a novel translational initiation mechanism. Our results show that SD strength is not a direct hallmark of translational efficiency in all bacteria. Instead, *Listeria* has evolved additional mechanisms to control gene expression level that are distinct from those utilised by *Salmonella* and *E. coli*.

*‘For the purpose of open access, the author has applied a Creative Commons Attribution (CC BY) licence to any Author Accepted Manuscript version arising ’*

## Introduction

Global transcriptome quantification techniques (e.g. RNASeq) are powerful methods to study the regulation of physiology and pathogenesis of *Salmonella*^40–45^. These studies have revealed that, despite the close linkage between prokaryotic transcription and translation, transcript levels often do not correlate with protein abundances^38, 46–51^.

Studies on translational control in bacteria have predominantly been based on the Gram-negative bacterium *E. coli* as a model organism and have shown that a combination of Shine-Dalgarno (SD) strength and optimal codon usage amongst other factors contribute to efficient translation^1–3^. Despite these important studies, translational control in other Gram-negative bacteria differs greatly, for example some phyla such as *Aquificota* and *Bacteroidota* naturally lack SD sequences^4–6^. Gram-positive bacteria are even more evolutionarily distant, and their translational control is poorly understood. Given that a significant number of infection-related deaths in humans are associated with bacterial pathogens (both resistant and susceptible to antimicrobials)^7^, it is crucial that we have a better understanding of bacterial translational regulation. We therefore investigated two representative bacterial pathogens – the Gram-negative *Salmonella enterica* serovar Typhimurium (*S*. Typhimurium) and the Gram-positive *Listeria monocytogenes* (*L. monocytogenes*), to understand the global translation landscape and its relationship with transcriptional control, particularly for the virulence machinery.

Non-typhoidal serovars of *Salmonella* are the leading cause of food-borne gastroenteritis; they infect millions and cause *c.* 230,000 fatalities each year^8^. *S.* Typhimurium is the most common non-typhoidal *Salmonella* strain isolated from patients around the world and is used in mouse models to study bacterial pathogenesis and host-microbe interactions by Gram-negative pathogens^9, 10^. In addition to infecting humans, *S.* Typhimurium is an important pathogen in livestock including chickens, pigs and cattle^11, 12^. It utilises a multitude of virulence factors to reach and invade host cells, and to support its intracellular survival^13–16^.

Most *Salmonella* virulence factors are encoded within horizontally acquired genomic regions known as *Salmonella* pathogenicity islands (SPIs)^17–21^. SPI-1 encodes a type III secretion system (T3SS), also known as an injectisome, which is responsible for the trigger-based invasion mechanism utilised also by many other Gram-negative bacteria. The SPI-1 T3SS penetrates host cells and secretes effector proteins through the needle to facilitate invasion of intestinal cells^19, 20, 22–24^. A further four pathogenicity islands (SPI-2, 3, 4 and 5) encode virulence factors required for infection and survival within host cells^20, 21, 25^. In addition to SPI-encoded virulence factors, some virulence factors are encoded outside of SPIs. Key examples are genes which encode additional effector proteins (e.g. SopA and SopE), which promote bacterial entry into host cells, regulate bacterial survival within host cells and control inflammatory responses^22, 26, 27^. Another key virulence factor is the bacterial flagellum, a large macromolecular rotary motor that enables cell motility, including to sites of infection^28–30^. A major component of the flagellum is the protein, flagellin, an important antigen which triggers a range of immune responses in host cells^31–35^.

The Gram-positive bacterium *Listeria monocytogenes*, the leading cause of listeriosis, is one of the most virulent foodborne pathogens, with a high rate of death associated with infection. Like *Salmonella*, *Listeria* encodes a range of virulence factors to promote entry and survival within host cells. Upon invasion of host cells, these bacteria replicate within the phagosome and produce listerolysin O (LLO), which lyses the phagolysosomal membrane, allowing the bacteria to escape into the cytoplasm, where they proliferate. They use cell-surface virulence factors to promote host actin polymerisation and thus mediate actin-based motility within the cytoplasm and into neighbouring cells, allowing dissemination within host tissues. Interestingly, listeriolysin O levels are translationally regulated to control virulence, highlighting the role of this control mechanism on bacterial pathogenesis^36^.

To investigate translational control in *Salmonella* and *Listeria*, we utilised ribosome profiling (RiboSeq) which involves deep sequencing of ribosome protected mRNA fragments to directly capture protein synthesis in a natural setting. Initially developed in yeast, RiboSeq is a highly sensitive global method that reveals the translatome at the time of harvest^37^. The technique determines the positions of ribosomes by exploiting the protection of a discrete fragment of mRNA (∼30 nucleotides) from nuclease digestion conferred by a translating ribosome^37, 38^. Deep sequencing of these ribosome-protected fragments (RPFs) generates a high-resolution view of the location and abundance of translating ribosomes on different mRNA species. The combination of RNASeq and RiboSeq datasets provides a global picture of: (1) mRNA abundance; (2) total protein synthesis (i.e. total number of ribosome-protected fragments on all mRNAs per gene – a direct measurement of the total amount of proteins being synthesised at the time of harvest); and (3) translation efficiency, a measurement of how well each mRNA is being translated (i.e. number of ribosome-protected fragments per mRNA). However, obtaining high quality RiboSeq datasets from bacteria has been problematic due to numerous technical difficulties and, so far, there are limited wild-type bacterial ribosome profiling data of sufficient resolution to reflect the triplet periodicity of decoding, which is instrumental to accurately identify short ORFs and non-canonical control mechanisms^39^. Here, we optimised RiboSeq for both model intracellular pathogens *Salmonella enterica* and *Listeria monocytogenes*. Our data display precise triplet phasing and, for the first time, allow us to determined their global translatome, permitting an accurate global characterisation of bacterial translational control.

### Measurement of the steady-state translatomes of *Salmonella* and *Listeria* with high-definition RiboSeq

To dissect the regulatory layers of virulence and pathogenicity of *Salmonella* and *Listeria*, we generated highly phased RiboSeq and parallel RNASeq data (Fig. 1A, S1, see extended material), allowing us to accurately uncouple RNA abundance from translation efficiency of cellular components directly relevant to virulence. The total translation (protein synthesis) of a given gene is dependent on both its mRNA abundance and the efficiency with which it is translated. As we were interested in components immediately relevant to infection and virulence, cells were harvested at OD_600_ of 1, when peak production of the SPI-1 transcriptional master regulator, HilA, occurs in *Salmonella*, and at mid-log growth phase for *Listeria*^52, 53^. These conditions allowed us to capture translation within *Salmonella* and *Listeria* that are ‘primed’ for infection^54^.

**Figure 1.**
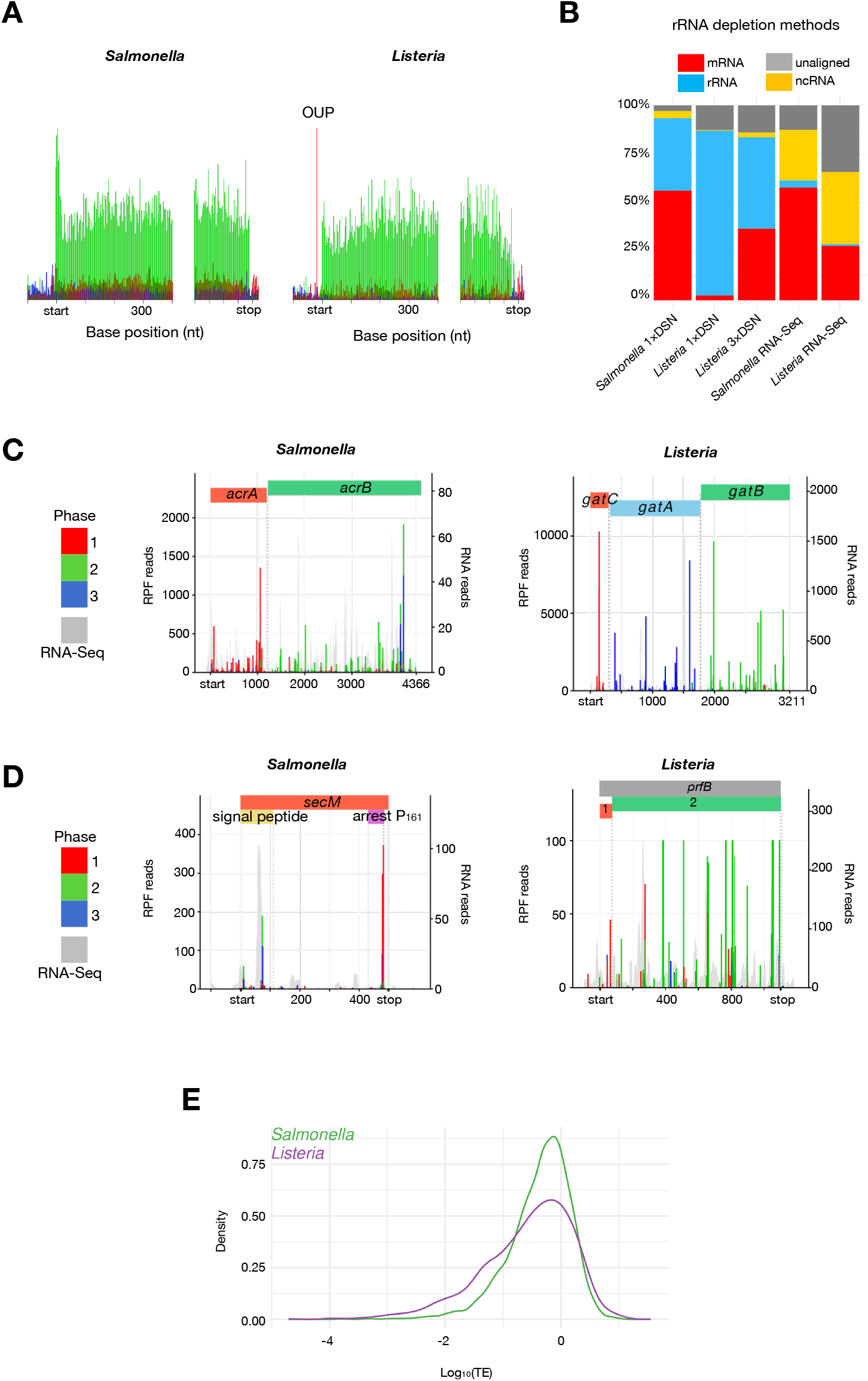
Tandem-RNASeq and RiboSeq of *Salmonella* and *Listeria* cells. **A**. Metagene translatome plots generated by aligning all coding sequences using the start and stop codons as anchors. Reads from the *Salmonella* SL1344 RiboSeq library (left) or *Listeria* 10403S RiboSeq library (right) that map to codon positions 1, 2 and 3 are coloured in red, green and blue, respectively. The plot was produced with the R software package riboSeqR^56^. **B**. Library composition of RiboSeq libraries from *Salmonella* (SL1344) or *Listeria* (10403S) generated with RNase I that were subjected to oligonucleotide-based rRNA subtraction, followed by one or three rounds of DSN treatment. **C**. The composition of RNASeq libraries post oligonucleotide-based rRNA subtraction is on the right. Translation visualisation of multi-cistronic transcripts: the *Salmonella acrA*-*acrB* operon, the *Listeria gatC*-*gatA-gatB* operon; **D**. the *Salmonella secM* gene which utilises programmed ribosomal pausing; and the *Listeria prfB* gene that utilises -1 programmed ribosome frameshifting. Red, green and blue bars indicate RiboSeq reads mapping to phases 1, 2 and 3, respectively, relative to the first nucleotide of the transcript. Grey shaded peaks show parallel RNASeq data. **E**. Cumulative distribution of translation efficiency for all *Salmonella* (green) and *Listeria* (purple) genes.

In addition, we also addressed a major obstacle in bacterial ribosome profiling: the removal of rRNA species, which account for a significant proportion of the total reads (> 95%), restricting analysis to highly abundant transcripts unless immense resources are dedicated to sequencing^55^. We therefore utilised a combination of rRNA subtraction using subtraction kits based on oligonucleotides anti-sense to bacterial rRNA, and duplex-specific nuclease (DSN)-based depletion treatment, to substantially reduce the proportion of rRNA reads within libraries, thereby substantially enriching libraries for reads corresponding to ribosome-protected fragments of mRNAs^55, 56^ (Fig. 1B). While the combination of anti-sense rRNA-based subtraction and one round of DSN treatment was effective at depleting reads corresponding to non-coding RNAs (ncRNA) including rRNA from *Salmonella* RiboSeq libraries, *Listeria* RiboSeq libraries required two further rounds of DSN treatment to sufficiently enrich for reads corresponding to RPFs (Fig. 1B).

Upon obtaining substantial sequencing depth, ∼97% (4543/4682) of *Salmonella* genes and ∼99% (2768/2800) of *Listeria* genes passed our filtering criteria for downstream analysis (see extended material). Quality control analysis was performed with particular attention to commonly known artefacts in bacterial RiboSeq, which could lead to over-interpretation of data^57^. First, artefacts in bacterial RiboSeq studies can arise due to the choice of nuclease used to generate RPFs^48, 58^. RNase I is typically used to generate RPFs from eukaryotes, whereas its use in *E. coli* has been less successful, reportedly due to its inhibition by *E. coli* ribosomes^59^. For this reason, bacterial RiboSeq studies tend to use S7 micrococcal nuclease (MNase). However, S7 MNase exhibits significant sequence specificity, resulting in RiboSeq data that contains high levels of noise and a lack of triplet phasing^57^. We titrated S7 MNase and RNase I to determine the optimal nuclease concentration to generate RPFs (Fig. S2). Metagene translatome analysis revealed accumulation of reads corresponding to translation initiation, regardless of nuclease treatment (Fig. 1A, S2B). Similar to previous reports, we found that treatment with S7 MNase resulted in data that do not reflect triplet phasing^47, 48, 57, 58^ (Fig. S2). In contrary to previous work in *E. coli*, treatment with RNase I, however, resulted in RPFs with a distinct size distribution that are highly phased, indicating that the enzyme can be used to generate *Salmonella* and *Listeria* RiboSeq data with single nucleotide resolution, visible at the individual gene level (Fig. 1A, 1D, S2, S3). Further, despite, a correlation of protein synthesis levels of genes in libraries generated with S7 MNase and RNase I, it was apparent that S7 MNase-treated RiboSeq datasets tended to over-estimate protein synthesis, especially for genes which are poorly translated (Fig. S2C).

**Figure 2.**
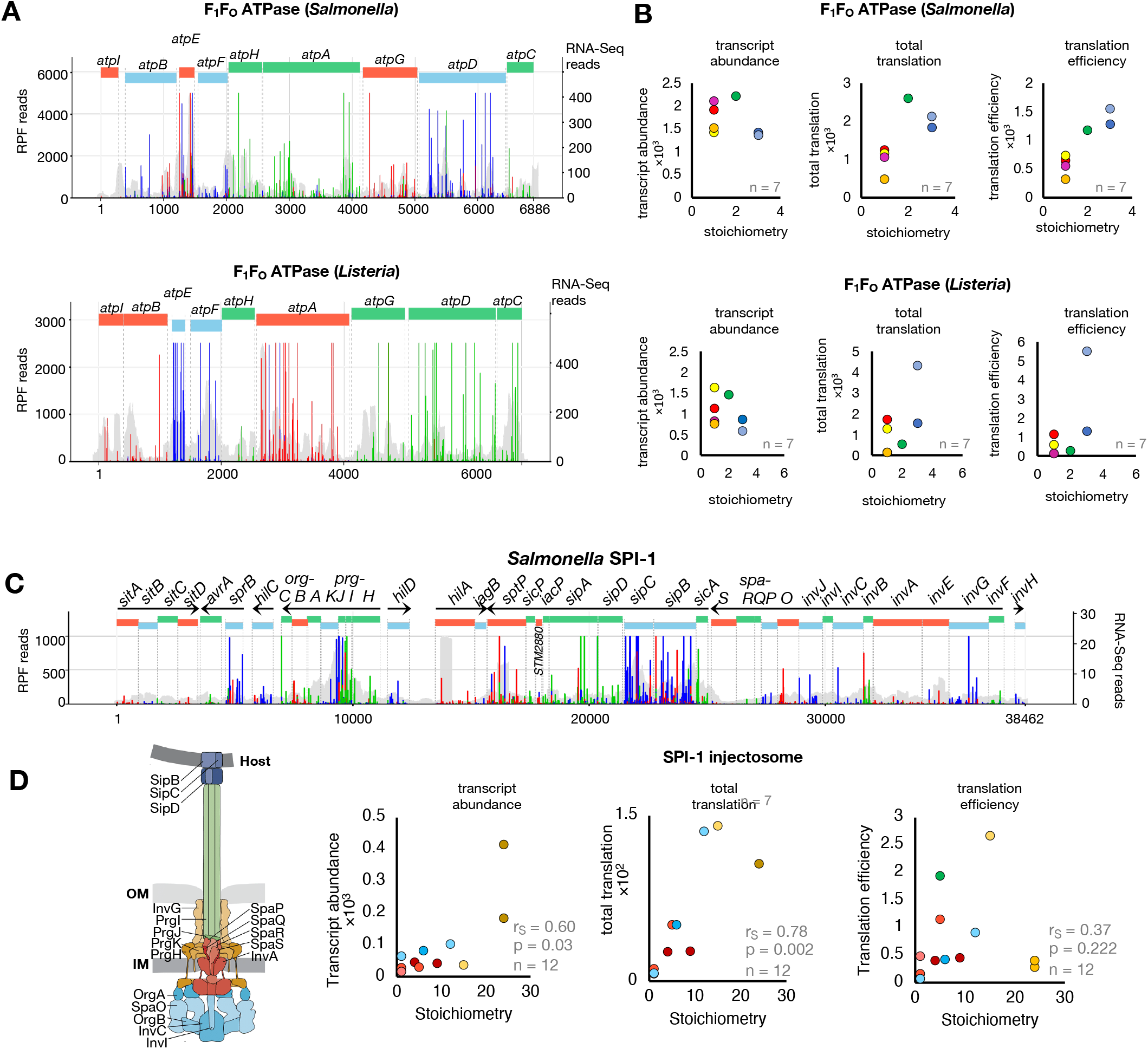
Proportional synthesis of protein complexes associated with virulence. **A**. Translation visualization *of Salmonella* and *Listeria atp* operon which contains the genes encoding components of the F_1_F_o_-ATPase complex (top and bottom, respectively). Red, green and blue bars indicate RiboSeq reads mapping to frames 1, 2 and 3, respectively. Grey shaded peaks show parallel RNASeq data. RiboSeq axis adjusted to enable visualisation of less translated ORFs **B.** Relationship between stoichiometry of F_1_F_0_ ATPase complex subunits and the corresponding transcript abundance, protein synthesis or translation efficiency. The data point corresponding to the *atpE* gene is an outlier due to significant ligation bias and was excluded from correlation analyses. **C**. Translation visualization of the *Salmonella* SPI-1 regulon which is composed of 8 transcripts (annotated with black arrows where arrow heads indicate the direction of transcription) and codes for virulence genes, including structural components of the SPI-1 injectosome. Red, green and blue bars indicate RiboSeq reads mapping to frames 1, 2 and 3, respectively. Grey shaded peaks show parallel RNASeq data. **D**. Relationship between stoichiometry of SPI-1 vT3SS structural genes and the corresponding transcript abundance, protein synthesis or translation efficiency (Right). Schematic representation of the SPI-1 vT3SS highlighting structural proteins (Left). Schematic modified from Wagner et al., 2018^13^.

**Figure 3.**
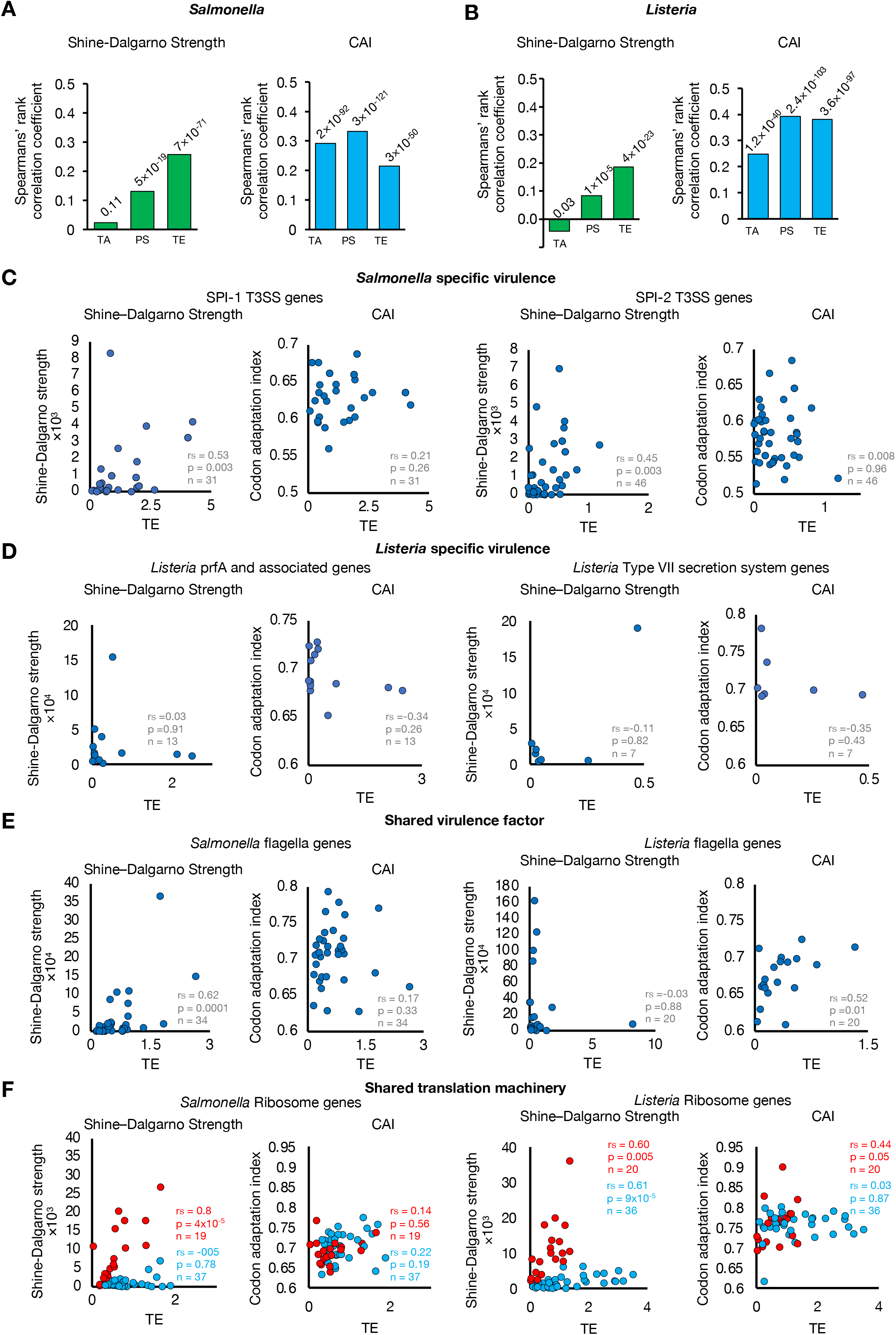
Regulation of proportional protein synthesis in *Salmonella* and *Listeria*. **A and B**. Histograms showing the Spearman’s rank correlation coefficients for the relationship between SD strength (left) or codon adaptation index (CAI, right) with transcript abundance (TA), protein synthesis (PS) or translation efficiency (TE) of all *Salmonella* and *Listeria* genes, respectively. **C**. The approximate p values have been calculated and are displayed above each histogram. SD strength (left) or codon adaptation index (CAI, right) plotted as a function of translation efficiency for *Salmonella* specific virulence factors: *Salmonella* SPI-1 T3SS structural genes (left) or *Salmonella* SPI-2 regulon genes (right); **D. Listeria specific virulence factors**: *Listeria prfA* and associated genes (left) or *Listeria* type VII secretion system genes (right)**; E. Shared virulence factor:** *Salmonella* (left) and *Listeria* (right) flagella genes; **F. Shared translation machinery**: *Salmonella* (left) and *Listeria* (right) flagella ribosomal protein genes. Genes groups with distinct correlation between translation efficiency and SD strength are coloured in red or blue.

We then focused on in-frame reads, i.e. reads that genuinely reflects ribosome occupation, for all downstream analysis, complemented by parallel RNASeq to tease apart RNA abundance (i.e. total RNASeq) from protein synthesis (i.e. total RiboSeq). We processed the data in a similar manner to our previous studies^56, 60, 61^, followed by correlation of protein synthesis with RNA abundance. As expected, protein synthesis positively correlated with RNA abundance. Importantly, we also observed that a large number of mRNAs were synthesised but not translated (Fig. S2C, S2F, S2H). This supports the notion that RNASeq alone does not accurately reflect global protein levels in bacteria despite close coupling of transcription and translation^49^.

We were also able to identify distinct features of *Salmonella* and *Listeria* translation. First, while the quality of *Salmonella* and *Listeria* RiboSeq libraries are comparable (Fig. 1A, S2, S3), a subset of genes in *Listeria* had a large number of ribosomal footprints where the expected P-site maps to a position seven nucleotides upstream of the start codon; we termed these ‘out-of-frame upstream peaks’ (OUP, Fig. 1A, right). These OUP reads are absent in all *Salmonella* RiboSeq libraries. Sequence enrichment analysis revealed that *Listeria* genes containing the OUP on average possess a slightly stronger SD sequence than *Listeria* genes without the OUP (Fig. 1B). Secondly, we observed that RPFs of highly abundant read sizes (24, 25, 28 and 29 nucleotides) in *Salmonella* are more phased than other read sizes (Fig. S3), similar to previous reports of plant and algae (ribosome profiles^56, 60, 62, 63^. This observation was consistent with a library generated with a different *Salmonella* strain as well as *Listeria* on a separate occasion (Fig. S3)^56^, suggesting that the accessibility to nucleases of the mRNA exit channel of *Salmonella* and *Listeria* ribosomes is similar to that of *Arabidopsis* and *Chlamydomonas*, where they are less protected by ribosomal proteins than for mouse and human ribosomes. In contrast to the RiboSeq libraries, reads in our RNA-Seq datasets have the expected features, including broader read-size distributions, are not phased, and are equally enriched in the untranslated regions and coding regions of mRNAs^37, 55, 56, 60, 61, 64^ (Fig. S2, S3, S4).

**Figure 4.**
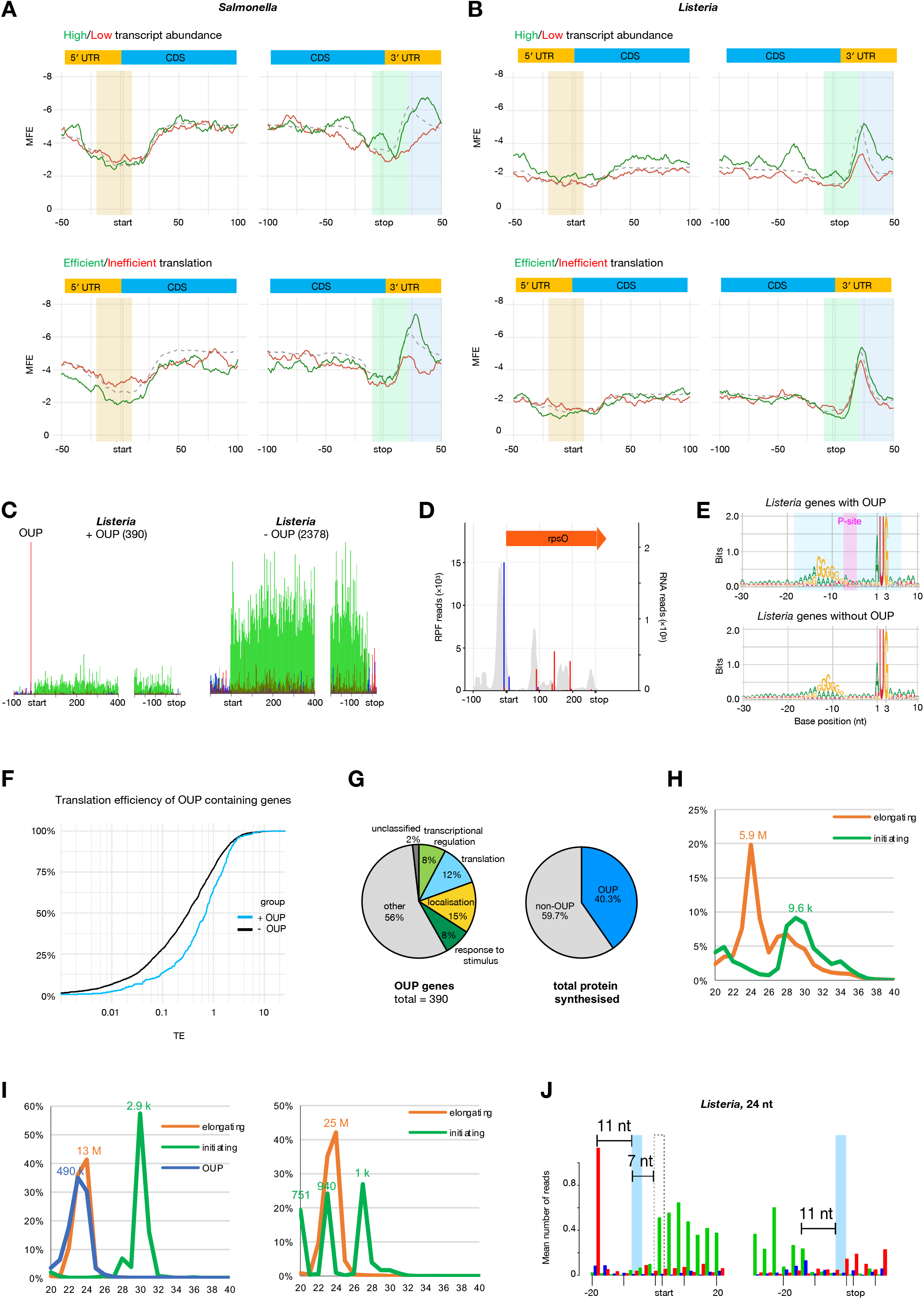
RNA secondary structure and OUPs regulating translation efficiency of genes in *Salmonella* and *Listeria*. **A and B.** Sliding-window average MFE of predicted secondary structure at each nucleotide of sequences surrounding the start and stop codons for the 100 most abundant (green) or least abundant (red) transcripts (top) or the 100 most efficiently translated (green) or least efficiently translated (red) transcripts (bottom) in *Salmonella* (A) or *Listeria* (B). Secondary structure in the highlighted areas is further compared in Fig. S5 and S6. **C**. Metagene translatome plots generated by aligning all open reading frames of Listeria containing the abundant out-of-frame upstream peak (OUP) of at least ten reads (left) or without (right). Reads that map to codon positions 1, 2 and 3 are coloured in red, green and blue, respectively. **D.** Translation visualisation of an *Listeria* OUP-gene *rpsO*. Red, green and blue bars indicate RiboSeq reads mapping to frames 1, 2 and 3, respectively. Grey shaded peaks show parallel RNASeq data. **E**. Nucleotide sequence logos for all *Listeria* genes with an upstream out-of-frame peak (OUP) of at least ten reads (top) and for all *Listeria* genes without OUP (bottom). The OUP footprint is highlighted in blue whilst the P-site is highlighted in magneta. **F**. The cumulative distribution for the translation efficiency of OUP genes (blue) and non-OUP genes (black). **G**. Pie chart of gene ontology distribution for OUP-genes (left). The ‘response to stimulus’ group excludes genes that also belongs to ‘transcriptional regulation’, ‘translation’ and localisation’. Overlaps are illustrated in Fig. S16E. Pie chart on right reflects proportion of all protein synthesised from OUP and non-OUP genes. **H**. Read size distribution from *Listeria* RiboSeq libraries for initiating ribosomes (green) or elongating ribosomes (orange). **I**. Read size distribution from *Listeria* RiboSeq libraries for initiating ribosomes (green), elongating ribosomes (orange) or OUP genes at the initiation site (blue). **J**. Read size distribution from *Salmonella* RiboSeq libraries for initiating ribosomes (green) or elongating ribosomes (orange).

The phased *Salmonella* and *Listeria* RiboSeq data also enabled direct visualisation of intrinsic features of bacterial translation, as illustrated by the following examples. First, the reading frames of individual genes were clearly visible, as exemplified by the *acrAB* operon in *Salmonella* (Fig. 1C, left) and the *gatABC* operon in *Listeria* (Fig. 1C, middle left). Secondly, we could detect ribosome pausing events such as following the ribosome arrest motif in *Salmonella secM* (Fig. 1D, middle right). Thirdly, we could detect changes in ribosome density at the known sites of programmed ribosomal frameshifting within the *Salmonella prfB* and *dnaX* genes and the *Listeria prfB* gene^65–69^ (Fig. S5, Fig. 1D, right).

Thus the translation of previously unannotated ORFs in *Salmonella* and *Listeria* is detected with high confidence (Table 1, Fig. S6). Finally, we revealed that global distribution of translation efficiency of *Salmonella* is more compressed than *Listeria* (Fig. 1E).

### Stoichiometric production through coordinated translation of protein complexes associated with virulence

We next assessed whether protein synthesis levels correlate with stoichiometry of components in multiprotein complexes. Components within multiprotein complexes are typically thought to be produced at levels which match their stoichiometry, as the excess production of one or more components over another could result in incorrect assembly, misfolding or aggregation, besides resource wastage^71, 72^. Li *et al*. (2014) showed that ∼55% of components in multiprotein complexes in *E. coli* are produced at levels indistinguishable from their stoichiometry ^47^. In *Salmonella* too, overall protein copy number positively correlates with translation efficiency, supported by the stronger correlation with protein synthesis than RNA abundance, suggesting that correct stoichiometry for a significant subset of proteins is determined translationally (Fig. S7, table 2).

We observed many of structural-component-related operons where protein copy number is directly controlled during translation (i.e. via translation efficiency). We therefore further investigated how these operons – many of which are associated with pathogenicity – are translationally controlled, starting with ATPase complexes. We first looked at the F_1_F_o_ ATPase complex, which in *Salmonella* is encoded by the *atp* operon. It consists of 9 genes, 8 of which encode subunits that assemble to form the F_1_F_o_ ATPase complex (Fig. 2A and 2B). The F_1_F_o_ ATPase is expressed from a single polycistronic transcript and should require individual genes within the operon to exhibit different translational efficiencies to reflect the stoichiometry of the complex^47^. *atpE* was removed from these analyses due to an overwhelming accumulation in read counts due to ligation bias in both the RNAseq and RiboSeq libraries, (Fig. S8). Similar to *E. coli,* we also observed a positive correlation between stoichiometry and translation efficiency in both *Salmonella* and *Listeria*, indicating that utilisation of translational control as a mechanism for proportional protein synthesis and stoichiometric assembly of the F_1_F_o_ ATPase complex is a common feature of both Gram-positive and Gram-negative bacteria^47^ (Fig. 2B). We also found that stoichiometric production of components of the multi-drug efflux components AcrAB and the iron-sulfur assembly components SufBCD are primarily regulated at the level of translation (Fig. S9A– C, S10). In contrast, stoichiometric production of components of the virulence T3SS ATPase complex is regulated by both transcript abundance and translation efficiency (Fig. S9D).

To better understand expression strategies utilised for the production of virulence factors, we assessed whether the stoichiometry of components within multiprotein complexes associated with virulence correlates with RNA abundance (transcription or RNA turnover) or translation efficiency and assessed whether the correlation is supported by the functions of individual components (e.g. whether T3SS effectors are expressed in excess of the injectisome structure through which they are secreted). We first examined the *Salmonella* SPI-1 locus, which, unlike the F_1_F_o_ ATPase, is composed of multiple operons, and we focused specifically on the structural components that assemble to form the SPI-1 injectisome type III secretion system (vT3SS) – a large multiprotein complex of known stoichiometry, ranging from one subunit copy to over 100 copies, that is essential for active host invasion (Fig. 2C-D). In contrast to F_1_F_o_ ATPase, stoichiometry for the majority of the T3SS structural components correlates well with transcript abundance, likely due to the expression derived from eight polycistronic transcripts, but more strongly with total protein synthesis, thus positive correlation with TE, suggesting that stoichiometry of SPI-1 structural components is co-transcription and translationally regulated (Fig. 2C-D).

### Regulation of proportional protein synthesis in *Salmonella* and *Listeria*

As our ribosome profiling revealed that the stoichiometry of many subunits within protein complexes in *Salmonella* and *Listeria* is translationally controlled, we reasoned that translation efficiency could be controlled by a combination of one or more factors. One is the strength of SD sequence, which provides an indication of initiation efficiency and is calculated through a combination of base-paring between a SD sequence and the anti-SD sequence on the 16S rRNA of the 30S ribosomal subunit, as well as the minimal free energy of local secondary structure surrounding the SD sequence and start codon which can interfere with 30S accessibility. The second factor is codon usage, which may affect regulation during elongation, and a third is secondary structure throughout the mRNA. To investigate these factors, we first determined the codon adaptation index (CAI) and SD strength of all translated *Salmonella* and *Listeria* genes and compared either CAI or SD strength score with transcript abundance, protein synthesis, or translation efficiency (Fig. 3A, 3B, S12, S13). Consistent with efficient initiation being the limiting factor for protein synthesis, the correlation between translation efficiency and SD strength is stronger in both *Salmonella* and *Listeria* than for transcript abundance or protein synthesis. This was particularly marked in the case of *Salmonella*. In contrast, correlation between CAI and translation efficiency is stronger in *Listeria* compared to *Salmonella.* As expected, little to no correlation was seen between transcript abundance and SD strength for either *Salmonella* or *Listeria* (Fig. 3A, 3B).

We next restricted the analysis to just *Salmonella* and *Listeria* virulence genes that encode components of multiprotein complexes (Figure 3C–F, S8-S11). For the SPI T3SS genes of *Salmonella*, we observed a significant positive correlation between SD strength and translation efficiency for both SPI-1 and SPI-2 genes while correlation with CAI was poor, suggesting that regulation for virulence machinery is largely determined through regulating translational initiation efficiency (Fig. 3C). As for virulence genes of *Listeria*, we observed no correlation between translation efficiency and either SD strength or CAI for *Listeria prfA* and associated genes, which was expected as these genes are not part of a multiprotein complex, although we also did not observe correlations for *Listeria* Type VII secretion system genes (Fig. 3D).

We next focused on multiprotein complexes that exist in both *Salmonella* and *Listeria*, starting with F_1_F_o_ ATPase, *sufB-D* and flagella (Fig. 3G-H, Fig. S9-10, S14-15). While we generally observed positive correlation between SD strength and translation efficiency in *Salmonella*, we observed no correlation for either SD strength or CAI for any of these complexes in *Listeria*, with the exception of flagella genes where translation efficiency positively correlates with CAI (Fig. 3E). Finally, we examined one of the most evolutionarily conserved multiprotein complexes, the ribosome (Fig. 3F). In *Salmonella,* we identified two groups of ribosomal proteins where translation efficiency either strongly correlates with SD strength, or not at all (Fig. 3F, Fig. S15). In contrast, while there are also two groups of ribosomal proteins in *Listeria,* both groups display a positive correlation between SD strength and translation efficiency with one group being more sensitive to the SD strength (in blue, Fig. 3F and S16C). Further, the group of *Listeria* ribosomal proteins that are less sensitive to SD strength positively correlate with CAI (in red, Fig 3F, S16B). Overall, these results indicate that regulation of translation initiation in *Salmonella* is more directly influenced by SD strength compared to *Listeria*.

### *Cis-*regulatory elements that facilitate efficient translation in bacteria

SD strength is calculated through a combination of degree of base pairing between the SD sequence and the anti-SD sequence of the 16S rRNA as well as local RNA secondary structure. We therefore investigated the relationship between translation efficiency and both factors. First, we hypothesised that transcripts that are inefficiently translated have a preference to form stronger RNA secondary structure, which may impede accessibility of the SD sequence to the 16S rRNA^73–76^. To test this, we grouped genes into four groups, comprising the top 100 most or least abundant transcripts or efficiently translated genes (i.e. 2.2% and 3.6% of genes in *Salmonella* and *Listeria*, respectively). We then predicted the average minimum free energy (MFE) of RNA secondary structure throughout the transcript for each group of genes within a 30-nt-wide sliding window^60^ (Fig. 4A–B). These analyses revealed that *Salmonella* mRNAs that are efficiently translated tend to be less structured around the start codon compared to transcripts that are not efficiently translated, as predicted (Fig. 4A, S18). Interestingly, we also noted that weakly structured 3′ UTR is a distinct feature for *Salmonella* mRNAs that are low in abundance or inefficiently translated (Fig. 4A, S18). In contrast to *Salmonella*, *Listeria* transcripts are generally less structured throughout the 5’UTR and CDS, with mfe ≈ -2 until 3’UTR. In addition highly abundant *Listeria* transcripts tend to be more structured in the coding region and 3’UTR (Fig. 4B, S19). However, we only saw a marginally weaker structure between the start codon and ∼25 nt upstream of the start codon and no significant difference in RNA secondary structure throughout the rest of mRNA as a function of translation efficiency (Fig. 4B, S19), despite the difference between the 100 most and least efficiently translated *Listeria* transcripts being much greater than *Salmonella* (Fig. 1E).

### *Listeria* genes containing Out-of-frame Upstream Peaks (OUPs) are more efficiently translated

While there is a lack of global preference for weak 5’UTR secondary structure in efficiently translated *Listeria* transcripts, we observed a highly unusual feature in a subset of such genes in that they contain a high accumulation of out-of-frame 22-24 nt footprints where the P-site is inferred to lie seven nucleotides upstream of the start codon. These out-of-frame upstream peaks (OUPs) were observed in ∼15% of all translated genes and were only observed in *Listeria* RiboSeq libraries and not in *Salmonella* RiboSeq libraries (Fig. 1A, 4C and D, S3B, S4A). Nucleotide sequence logos revealed that *Listeria* genes containing an OUP contain on average a marginally stronger SD sequence and more often initiate with an AUG codon compared to non-OUP *Listeria* genes (Fig. 4E, 4F and S20A). In addition, the site of the OUP-footprint is not significantly less structured than in non-OUP genes (Fig. S20F). Importantly, comparison between OUP and non-OUP genes showed that mRNAs of OUP genes are more abundant and more efficiently translated than mRNAs of non-OUP genes (Fig. 4F). Moreover, OUP genes generally encode proteins involved in regulation of translation, transcription, response to stimulation or localisation/membrane transporters (Fig. 4G, Fig. S20D-E, and table 4). Despite comprising just 15% of genes (390/2768), OUP genes were responsible for 40% of all protein production (Fig. 4G).

To gain further insight into OUP-mediated translation, we next investigated the read size distribution of different classes of ribosome protected footprints, namely initiation (where inferred P-site aligns to the start codon) and elongation footprints in both *Salmonella* and *Listeria* and OUP in *Listeria* (Fig. 4H-J). As expected, the size distribution of *Salmonella* initiation and elongation footprints peaks at 28-30nt, and 24nt, respectively (Fig. 4H). The larger *Salmonella* initiation footprint is likely due to the presence of initiation factors 1 and 2 during initiation and/or alternative ribosome conformation present when the SD sequence interacts with the anti-SD sequence within the 30S subunit, together with positioning of the start codon at the P-site. Intriguingly, in *Listeria,* the size distribution of OUP footprints is ∼22-24 nt, similar to elongation footprints for both OUP and non-OUP genes, as well as a small proportion of non-OUP initiation footprints, while initiation footprints for OUP genes are significantly larger at ∼29-32 nt compared to initiation footprint for non-OUP genes (20, 23 and 27 nt) (Fig. 4I and J).

### The sequence context surrounding the start codon has a greater impact on *Salmonella* translation than for *Listeria*

To further investigate additional *cis-*regulatory elements that may contribute to translation efficiency, whether during initiation or at the early stages of elongation, we investigated sequences surrounding the start codons, both in terms of nucleotide and amino acid identity. Conservation of sequences upstream of and flanking start codons of the most efficiently translated genes in *Salmonella* revealed that the SD sequence has a preference for an adenosine followed by two tandem guanosine nucleotides at positions -10 to -8 compared to inefficiently translated genes (Fig. 5A). In contrast, efficiently translated *Listeria* genes have a preference for adenosine at position -3 and -4, a feature similar to the eukaryotic Kozak consensus sequence^77^, but only minor differences in SD sequence compared to genes that are inefficiently translated (Fig. 5B). This complements our analysis above where SD strength is a major determinant of translation efficiency in *Salmonella*. In *Listeria*, however, the less clear difference in nucleotide preference in the SD sequence as well as local secondary structure between efficiently and inefficiently translated genes complements our observation of poor correlation between SD strength, translation efficiency and stoichiometry (Fig. 3). Furthermore, our *Salmonella* data revealed that SD sequences of inefficiently translated transcripts tend to be further away from the start codon compared to those of efficiently translated transcripts (Fig. S20B), consistent with previous reports in *E. coli*, and supporting the notion of spacer length being an important modulator of initiation efficiency^2, 78, 79^. However, we did not observe differences in spacer length for *Listeria,* except perhaps for a small proportion of OUP-genes where the SD sequence is 1 nt further away (S20B-C). These observations highlight the distinct differences in the regulation of translation initiation between the two evolutionarily divergent bacteria.

**Figure 5.**
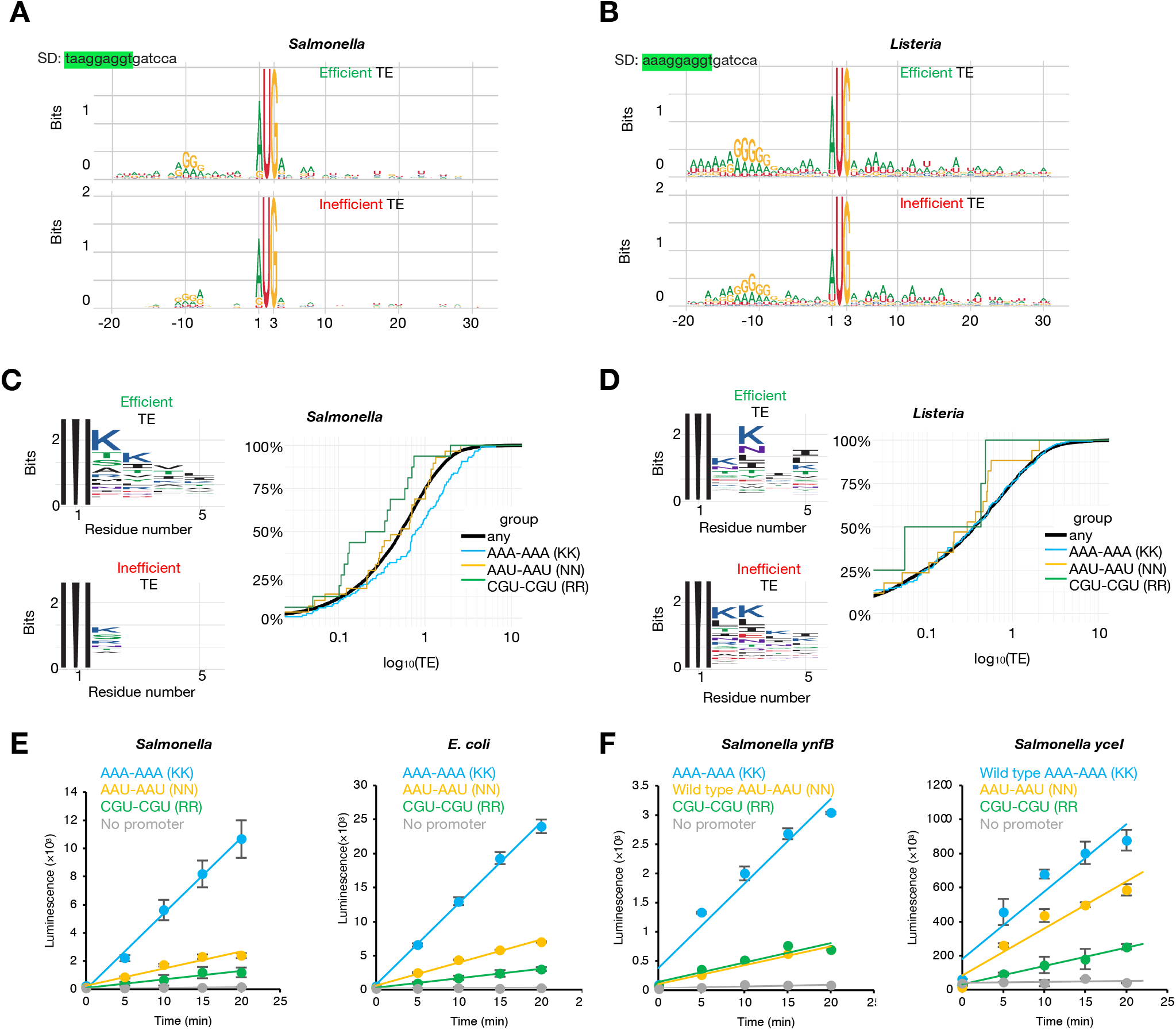
Sequence features regulating translation efficiency of genes in *Salmonella* and *Listeria*. **A and B**. Sequence logos of sequences surrounding the start (left) or stop codon (right) of the top 100 most efficiently translated genes (top) or least efficiently translated genes (bottom) in *Salmonella* (A) and *Listeria* (B), respectively. The SD sequence based on the sequence of the *Salmonella* or *Listeria* 16S rRNA sequence is highlighted in green (top left). **C and D**. Sequence logos of amino acid sequences surrounding the start codon of the top 100 most efficiently translated genes (top left) or least efficiently translated genes (bottom left) in *Salmonella* (A) and *Listeria* (B), respectively. The cumulative distribution of translation efficiency for genes containing AAA-AAA, AAU-AAU or CGU-CGU after the start codon is on the right. **E**. Induction time courses of LuxA luciferase expression measured by luminescence. Constructs containing AAA-AAA (KK, blue), AAU-AAU (NN, orange) or CGU-CGU (RR, green) or a construct lacking a promoter (grey) were expressed in *Salmonella* (SL1344) or *E. coli* (MC4000). **F**. Induction time courses of LuxA luciferase expression measured by luminescence where LuxA is fused with the first 30 nt of *ynfB* (which natively begins with AAU-AAU after the start codon) or the first 30 nt of *yceI* (which natively begins with AAA-AAA, after the start codon) to LuxA, or their mutants (as indicated).

Unexpectedly, we also detected an enrichment for adenosine at the 4^th^, 5^th^, 7^th^ and 8^th^ nucleotides in the coding region of efficiently translated genes in *Salmonella*, which would correspond to an enrichment of either asparagine (AAU, AAC) or lysine (AAA, AAG) residues at the 2nd and 3rd amino acid positions (Fig. 5A). A similar enrichment for adenosine nucleotides was observed in the coding region immediately following the start codon in *Listeria*, although there is similar enrichment also for inefficiently translated genes (Fig. 5B). Inspection of the amino acid sequences of genes that are efficiently translated revealed an enrichment for lysine residues over asparagine residues at both sites (Fig. 5C, 5D, S21). As lysine residues contain positively charged side chains, we assessed whether there is also a bias towards coding for positively charged arginine residues at the 2^nd^ and 3^rd^ codon positions in efficiently translated genes. To do this we stratified all *Salmonella* and *Listeria* genes based on whether they code for two tandem arginine, lysine or asparagine residues at the 2^nd^ and 3^rd^ codon positions. We found that in *Salmonella*, genes coding for two tandem lysine residues at the 2^nd^ and 3^rd^ positions were translated more efficiently than average and, interestingly, genes coding for two tandem arginine residues at the 2^nd^ and 3^rd^ positions were translated less efficiently than average, despite reported similar lysine and arginine tRNA abundances and both amino acids containing positively charged side chains^80^ (Fig. 5C). The translation efficiency of genes encoding two tandem asparagine codons was comparable to average. In contrast, we observed no significant difference in translation efficiency depending on the presence of two tandem lysine, arginine or asparagine codons at the 2^nd^ and 3^rd^ codon positions in *Listeria* (Fig. 5D, right).

To test whether the presence of lysine, arginine or asparagine codons at the 2^nd^ and 3^rd^ codon positions influences translation efficiency we utilised a luciferase (LuxAB) translational reporter assay where transcription is controlled by a tetracycline inducible promoter, in both *Salmonella* and the Gram-negative *E. coli* (Fig. 5E). Strikingly we found that the presence of two lysine codons (AAA-AAA) at the 5′ end of the *LuxA* coding region significantly increases reporter luminescence, which was not observed for two arginine (CGU-CGU) nor for two asparagine (AAU-AAU) codons (Fig. 5E). The luminescence increase for the di-lysine-codon-containing reporter was apparent shortly after transcriptional induction, and transcript levels remained similar between all constructs throughout the time course (Fig. S22).

To further investigate the effect on translation efficiency of di-lysine (AAA-AAA), di-asparagine (AAU-AAU) or di-arginine (CGU-CGU) at the 2^nd^ and 3^rd^ codon positions, we fused to *luxA* the first 10 codons of the wild-type *Salmonella* gene, *ynfB*, (which natively begins with AUG-AAU-AAU) or its variants in which the tandem AAU-AAU are replaced with AAA-AAA or CGU-CGU. Luciferase reporters containing the AAA-AAA replacement gave significantly higher luminescence than reporters containing the arginine replacement or the wildtype AAU-AAU sequence (Fig. 5F, left). Similarly, we fused to *luxA* the first 10 codons of the wild-type *Salmonella* gene, *yceI*, (which natively begins with AUG-AAA-AAA) or its variants in which the tandem AAA-AAA are replaced with AAU-AAU or CGU-CGU (Fig. 5F, right). Reporter constructs where the wild type AAA-AAA sequence is replaced with tandem AAU or CGU codons gave lower luminescence than reporters containing the wild type sequence. Overall, these results are in agreement with our RiboSeq data whereby presence of two lysine codons AAA-AAA at the 5’ end of ORFs correlates with efficient translation and show that this feature is functional in both *Salmonella* and *E. coli*.

## Discussion

Through accurate determination of translation efficiency of all expressed mRNAs in *Salmonella* and *Listeria*, we have identified multiple regulatory mechanisms that contribute towards efficient translation and virulence factor production. Moreover, we have shown that stoichiometric control of components of multiprotein complexes that drive virulence is hardwired within the messenger RNAs to directly control differential translation in *Salmonella* but not in *Listeria*.

We confirmed that a strong SD sequence as well as weak RNA secondary structure around the start codon are instrumental for efficient translation in *Salmonella,* presumably to enable efficient 30S subunit joining to the messenger RNA. In addition, we discovered that presence of AAA at the 2nd and 3rd codons of ORFs drives efficient translation in both *Salmonella* and *E. coli;* perhaps this sequence facilitates efficient transition of the 70S ribosome from initiation to elongation (Fig. 5G and Fig. 6). However, regulation of translation initiation differs substantially in *Listeria* (Fig. 5G). Besides greater variance than in *Salmonella*, average SD sequences between efficiently and inefficiently translated transcripts is almost indistinguishable, and there is also little difference in average secondary structure throughout the mRNA. However, this is likely a consequence of low GC% of the *Listeria* genome (37.8% compared to 52.2% in *Salmonella*), and therefore does not need to select against structure around the start codon to achieve the same lack of structure to promote efficient translation as in *Salmonella*.

**Figure 6.**
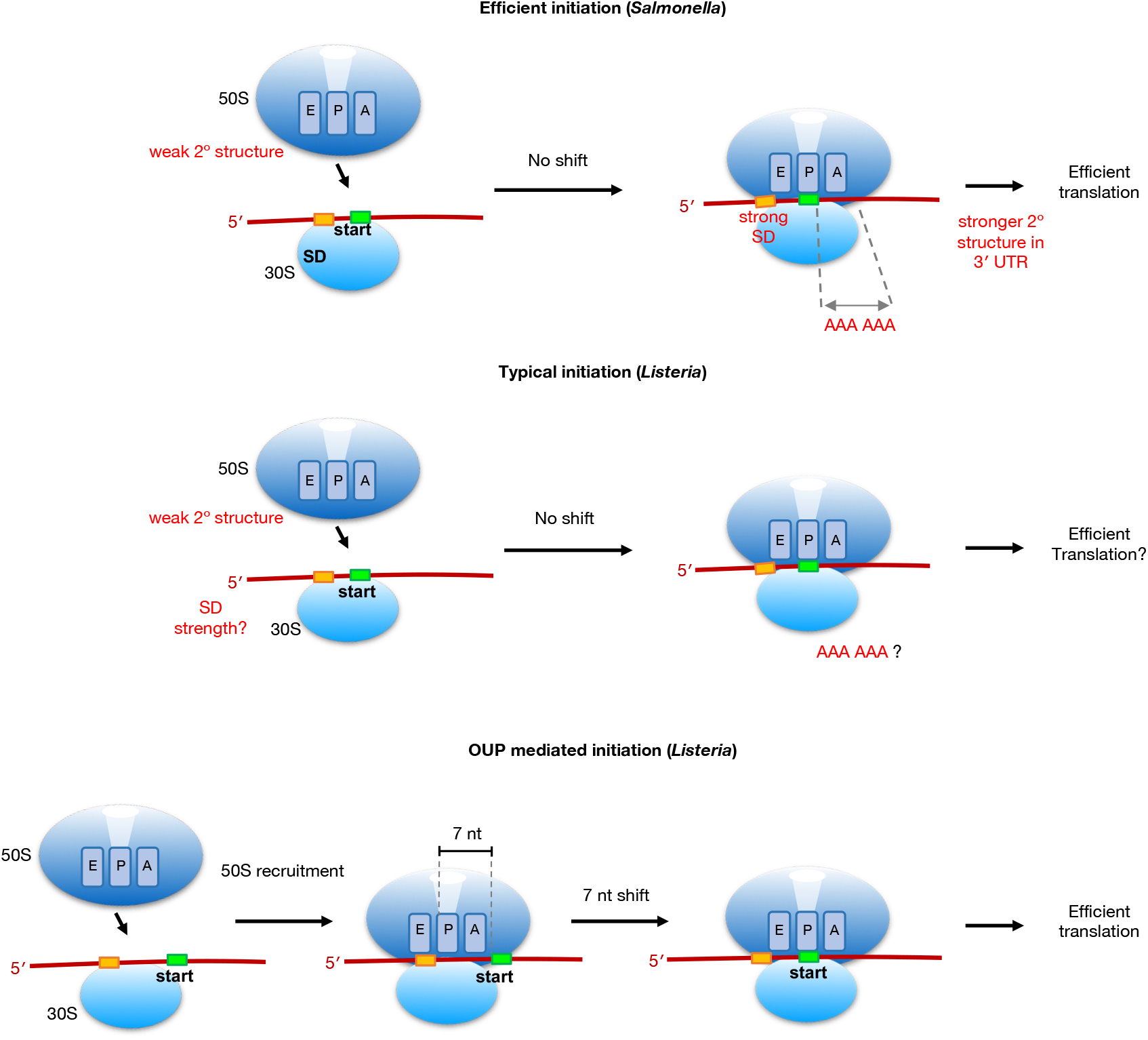
Models of *Salmonella* and *Listeria* translation initiation. Summary model showing *cis-*elements utilised for efficient translation in *Salmonella* (top) and *Listeria* (middle and bottom). In *Salmonella*, translation initiation is directly modulated by weak RNA secondary structure surrounding the start codon as well as interaction between the SD and anti-SD sequence. In addition, decoding of di-lysine codons AAA-AAA soon after initiation significantly enhances translation elongation. In contrast, in *Listeria,* there is no obvious preference for weak RNA secondary structure nor SD sequence nor di-lysine codons after the start codon for efficient translation. Instead, *Listeria* utilises two distinct translation initiation mechanisms, a canonical mechanism where the assembly of the ribosome occurs with the start codon at the P-site while a second distinct mechanism utilised by significant number of efficiently translated genes involves assembly of the ribosome 7 nt upstream of the start codon prior to migration of 7 nt until the P-site encounters the start codon for translational initiation (bottom).

Perhaps the most striking feature of efficiently translated *Listeria* genes are the OUPs, potentially indicating a novel mechanism of bacterial initiation. Canonical bacterial initiation begins with the 70S ribosome assembly with initiation factors 1 and 2, with the start codon positioned in the P-site, results in larger initiation footprint due to presence of initiation factors, as observed in *Salmonella* (Fig. S4D). It is possible that OUP footprints derive from a 30S subunit associated with initiation factors; however this is unlikely as pre-initiation 30S is thought not to sufficiently protect mRNA from nuclease digestion. Another possibility is that in *Listeria,* for OUP-mediated initiation 70S formation occurs further upstream in the absence of initiation factors 1 and 2, resulting in a similar footprint size as elongating ribosomes, followed by progression of 7 nt and acquisition of initiation factors and fMet-tRNA to initiate at the start codon, with a larger footprint size due to incorporation of initiation factors 1 and 2 (Fig. 6). A third possibility is that the ribosomes generating an OUP footprint have the start codon in the P-site, but this would necessitate structural rearrangements to account for the observed footprint size (22-24 nt). Additional factors and/or an unusual ribosome conformation could lead to the protection of 18 nt 5’ of the P-site rather than the normal 11 nt, and the conformation and/or empty A-site might make the mRNA 3’ end accessible to RNase1 cleavage closely downstream of the P-site AUG. Future work will investigate which of these hypothesis, if any, is correct.

Translational regulation is a critical process that governs the rate and efficiency of protein synthesis in all living cells. Our study has highlighted striking differences in translational regulation between two evolutionarily divergent pathogenic bacteria. The intrinsic molecular differences in translational regulation revealed by our analysis will provide new strategies for specific targeting of antibiotic-resistant bacteria, most importantly with respect to Gram-positive pathogens. Additionally, characterisation of these differences can lead to a better understanding of the basic molecular biology of these organisms, which can inform broader research in the field of microbiology.

## Materials and methods

Please see Bryant_Lastovka_et_el_2023_Extended_Material.xlsx

## Data availability

Raw and processed data are available from RiboSeq/RNASeq series (ArrayExpression accession E-MTAB-11527). Customised scripts used for this project are available upon request.

## Tables

See associated Bryant_Lastovka_et_el_2023_Tables.xlsx

## Conflict of interest

The authors declare no conflict of interest

## Acknowledgements

We would like to thank Dan Portnoy, University of Berkeley, for the generous gift of the *Listeria* 10403S strain, and Clare Bryant and Gillian Fraser, University of Cambridge, for the generous gift of the Salmonella SL1344 and SJW1103 LT2-derived strains, respectively. We would also like to thank Ian Brierley, John Carr, Julian Hibberd, Nancy Standart and Ben Luisi for discussions and comments on the manuscript. FL was supported by a BBSRC DTP studentship. BYWC and OJB are supported by a Medical Research Council Fellowship to BYWC [MR/R021821/1]. JP is supported by a BBSRC project grant awarded to BYWC [BB/V006096/1]. The BYWC laboratory is supported by a Medical Research Council Fellowship [MR/R021821/1] and BBSRC project grants [BB/X001261/1, BB/V017780/1 and BB/V006096/1] and a Royal Society Research Grant [RGS\R2\192222].

## Authors contribution

B.Y.W.C., O.J.B., F.L. conceived the research and designed experiments. B.Y.W.C., O.J.B. and J.P. generated RiboSeq and RNA-Seq libraries, O.J.B. performed most molecular experiments and *in vivo* functional assays. F.L. performed most of the bioinformatic analysis. B.Y.W.C., O.J.B. and F.L. wrote the manuscript.

---

**Figure S1.** A schematic illustrating our revised RiboSeq and RNA sequencing method. Translation was arrested by treating cells with highly concentrated chloramphenicol immediately prior to harvesting cells by rapid cooling, followed by centrifugation and flash freezing in liquid nitrogen for cryolysis. Clarified supernatants were used to generate RiboSeq and RNASeq libraries.

**Figure S2.** Read length distribution of RiboSeq libraries from *Salmonella* generated with S7 MNase (750 U or 1000 U) and corresponding RNAseq library (left) or generated with RNase I (500 U or 1000 U) (middle). The length distribution of RiboSeq libraries from *Salmonella* generated with RNase I (500 U) treated with RiboZero or with RiboZero in combination with DSN (right). **B**. Metagene translatome and metagene transcriptome plots generated by aligning all open reading frames of LT2-derived *Salmonella* using the start and stop codons as anchors. RiboSeq libraries were generated with S7 MNase I (500 U, left), RNase I (1000 U, middle). Reads that map to codon positions 1, 2 and 3 are coloured in red, green and blue, respectively. The plot was produced with the R software package riboSeqR^56^. **C**. Scatterplot showing the relationship between protein synthesis (i.e. RPF abundance) and transcript abundance (i.e. mRNA abundance) for all transcribed and translated *Salmonella* genes. **D**. Metagene transcriptome plot generated by aligning all open reading frames of pathogenic *Salmonella* SL1344. **E**. Scatterplot showing the relationship between total protein synthesis (i.e. RPF abundance) and transcript abundance (i.e. mRNA abundance) for all transcribed and translated *Salmonella* genes from strain SL1344. **F**. Metagene transcriptome plot generated by aligning all open reading frames of pathogenic *Listeria* 10403S. **G**. Scatterplot showing the relationship between protein synthesis (i.e. RPF abundance) and transcript abundance (i.e. mRNA abundance) for all transcribed and translated *Listeria* genes from strain 10403S.

**Figure S3. A.** RiboSeq and parallel RNASeq generated from LT2-derived *Salmonella* strain or pathogenic SL1344 across different RPF sizes. Red, green and blue bars indicate RiboSeq reads mapping to codon positions 1, 2 and 3, respectively. **B**. RiboSeq and parallel RNASeq generated from *Listeria* strain 10403S across different RPF sizes. Red, green and blue bars indicate RiboSeq reads mapping to codon positions 1, 2 and 3, respectively. **C and D**. Read size distribution of RiboSeq (left) and RNA-Seq (right) in *Salmonella* and *Listeria*.

**Figure S4. A.** Detailed metagene translatome plots of *Listeria* 10403S, *Salmonella* SL1344 and LT2-derived *Salmonella*. Reads that map to codon positions 1, 2 and 3 are coloured in red, green and blue, respectively. **B**. Scatterplot showing the relationship between protein synthesis (i.e. RPF abundance) and transcript abundance (i.e. mRNA abundance) for all transcribed and translated *Listeria* genes from strain 10403S. Genes containing the out-of-frame upstream peak (OUP) of at least ten reads are coloured in blue.

**Figure S5. Translation** visualisation of *Salmonella prfB* (top), *dnaX* (middle), *secM* and *secA* (bottom). Red, green and blue bars indicate RiboSeq reads mapping to codon positions 1, 2 and 3, respectively. Grey shaded peaks show parallel RNASeq data.

**Figure S6.** Translation visualisation of novel ORFs identified from the *Salmonella* RiboSeq data. Red, green and blue bars indicate RiboSeq reads mapping to codon positions 1, 2 and 3, respectively. Grey shaded peaks show parallel RNASeq data.

**Figure S7.** Transcript abundance (top), protein synthesis (middle) and translation efficiency (bottom) plotted against the published copy number of proteins reported in the literature^81–95^.

**Figure S8. A and B.** Relationship between stoichiometry of F_1_F_0_ ATPase complex subunits for *Salmonella* and *Listeria,* respectively and the corresponding transcript abundance, protein synthesis or translation efficiency. Far right panels present relationship of translation efficiency with SD strength and codon adaptive index, respectively. Data points corresponding to the *atpE* gene (blue) were outliers and included in the generation of these charts (but excluded from the equivalent charts in Figure 2).

**Figure S9. A.** Structure of the bacterial AcrAB-TolC drug efflux pump complex^96^. Transcript abundance, total protein synthesis and translation efficiency plotted as a function of the stoichiometry of the AcrA/AcrB components of *Salmonella*. **B**. Transcript abundance, total protein synthesis and translation efficiency of a representative range of bacterial drug efflux pump components of *Salmonella*. **C**. Correlation between transcript abundance, protein synthesis and translation efficiency plotted as a function of the stoichiometry of the SufB/SufC/SufD components of *Salmonella* (left). SD strength and codon adaptation index (CAI) plotted as a function of translation efficiency. **D**. Structural model of the *Salmonella* SPI-1 ATPase complex. Transcript abundance, total protein synthesis and translation efficiency plotted as a function of the stoichiometry of OrgB (green), InvC (blue) and InvI (red).

**Figure S10. A-C.** Transcript abundance, total protein synthesis and translation efficiency plotted as a function of the stoichiometry, and SD strength and codon adaptation index (CAI) plotted as a function of translation efficiency, for components of flagella (**A**), SufB-D (**B)** and components of F_1_F_O_-ATPase (**C**) genes.

**Figure S11.** Relationship between stoichiometry of SPI-1 vT3SS structural genes and the corresponding transcript abundance, protein synthesis or translation efficiency. PrgJ and PrgI are indicated on the chart.

**Figure S12.** Scatterplots showing the relationship between the codon adaptation index (CAI, left) or SD strength (right) and the translation efficiency of ribosomal protein genes in *Salmonella* and *Listeria*, respectively.

**Figure S13.** Scatterplots showing the relationship between the codon adaptation index (CAI, left) or SD strength (right) and the translation efficiency of *Listeria* ribosomal protein genes that are either OUP or non-OUP genes.

**Figure S14.** Scatterplots showing the relationship between the codon adaptation index and translation efficiency for SPI-1 T3SS and SPI-2 T3SS genes (**A-B**); ribosomal protein and flagella genes for *Salmonella* and *Listeria*, respectively (**C-F**); and *PrfA* and associated genes as well as Type VII secretion system genes in *Listeria* (**G-H)**. Spearman’s correlation coefficient and the approximate p value have been calculated.

**Figure S15** Scatterplot showing the relationship between the codon adaptation index and translation efficiency for all *Salmonella* ribosome genes, highlighting the two classes of genes (red or blue) (**A**) or the two classes of genes plotted separately (**B and C**).

**Figure S16** Scatterplot showing the relationship between the codon adaptation index with translation efficiency for all *Listeria* ribosome genes, highlighting the two classes of genes (red or blue) (**A**) or the two classes of genes plotted separately (**B and C**).

**Figure S17** Scatterplot showing the relationship between the codon adaptation index and translation efficiency for *Listeria* ribosome genes highlighting OUP genes (red) and non-OUP genes (blue).

**Figure S18.** Scatterplots showing the relationship between mRNA abundance, protein synthesis and translation efficiency and average MFE values for RNA secondary structure at the translation start site (left), translation stop site (middle) and in the 3′ UTR for all *Salmonella* genes. Spearman’s correlation coefficient and the approximate p value have been calculated. A summary of the Spearman’s rank coefficient for each plot is given at the bottom.

**Figure S19.** Scatterplots showing the relationship between mRNA abundance, protein synthesis and translation efficiency and average MFE values for RNA secondary structure at the translation start site (left), translation stop site (middle) and in the 3′ UTR for all *Listeria* genes. Spearman’s correlation coefficient and the approximate p value have been calculated. A summary of the Spearman’s rank coefficient for each plot is given at the bottom.

**Figure S20. A.** Nucleotide sequence logos generated from all *Listeria* genes containing an OUP of at least ten reads (left) or all other translated genes (right). **B.** Box and whisker plots showing distance in nucleotides between the SD sequence and the start codon for efficiently translated genes (top 100 genes by translation efficiency), inefficiently translated genes (bottom 100 genes by translation efficiency) and all genes (total) in *Salmonella* and **C**. *Listeria*. **C.** Histogram displaying the number of *Listeria* OUP (blue) or non-OUP (grey) genes that contain a specified distance between the SD and start codon. **D**. Bar chart showing the enrichment strength of and the number of genes that belong to a gene ontology group among *Listeria* OUP genes. These groups can overlap. **E**. Euler diagram showing the overlap of selected gene ontology groups in *Listeria* OUP genes. **F**. Sliding-window average MFE of predicted secondary structure at each nucleotide of sequences surrounding the start codon of OUP containing genes that contain at least 10 reads of the OUP (blue), genes not containing an OUP (black), efficiently translated genes (green, top 100 genes by TE) and inefficiently translated genes (pink, bottom 100 genes by TE). The OUP footprint is coloured in pale blue whilst the P-site is highlighted in magenta.

**Figure S21. A**. Sequence logos of N-terminal amino acid sequences of the top 100 most efficiently translated genes (top) or the top 100 least efficiently translated genes (bottom) in *Salmonella* (left) and *Listeria* (right). **B**. Venn of genes contains following features: weak secondary structure at the translation start site, strong secondary structure in the 3′ untranslated region, high SD strength, or AAA-AAA codons following the start codon. **C**. Pie chart showing the proportion of *Listeria* genes that either contain an OUP or do not (left). **D**. Pie chart of gene ontology distribution for non-OUP-genes. The ‘response to stimulus’ group excludes genes that also belong to ‘transcriptional regulation’, ‘translation’ and localisation’.

**Figure S22. A**. Transcript levels from induction time courses of luciferase expression from constructs containing N-terminal fusions of methionine-lysine-lysine (MKK, blue), methionine-asparagine-asparagine (MNN, yellow), methionine-arginine-arginine (MRR, green), or a construct lacking a promoter (P-less, grey) in *Salmonella* SL1344. The relative transcript abundance was determined by RT-qPCR with *lppA* (SL1344_1311) as a reference housekeeping gene and MKK at time point 0 as the reference sample. Three biological replicates were performed. The ribbons represent the standard error of the mean relative transcript abundance.

## Notes

### Competing Interest Statement

The authors have declared no competing interest.

### Summary of Updates

Modified minor error in schematic.

