## Supplementary Figures for "The distinct translational landscapes of Gram-positive and Gram-negative bacteria"

### Figure S1

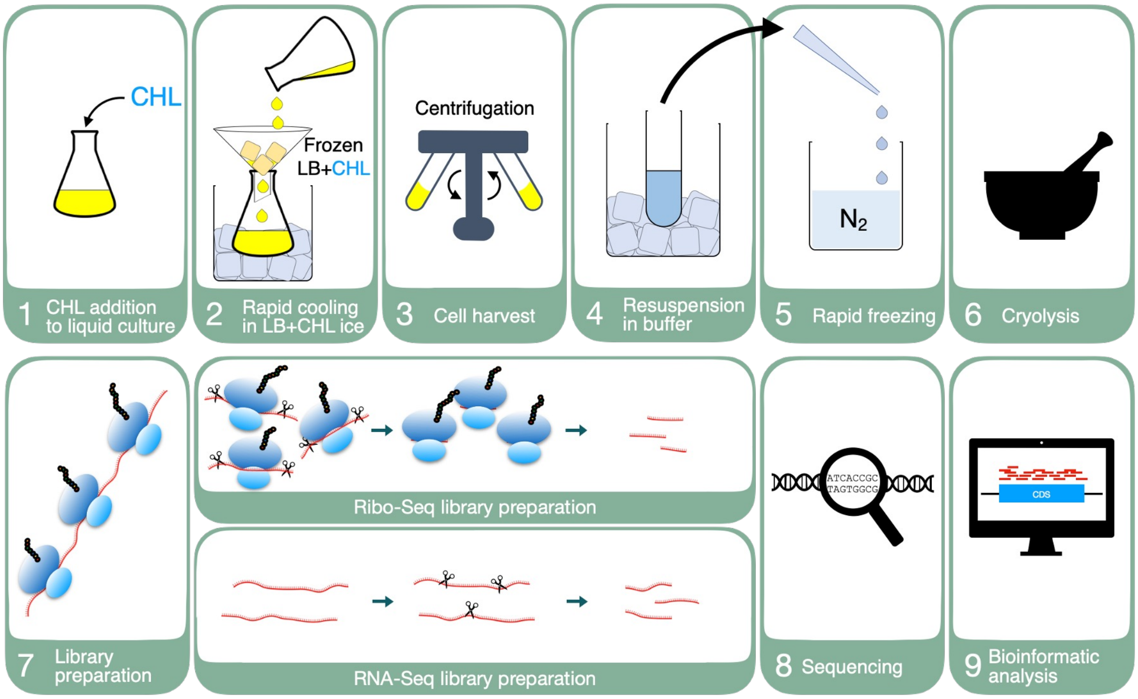

CHL
Frozen
LB+CHL
Centrifugation
$N_2$
1 CHL addition to liquid culture
2 Rapid cooling in LB+CHL ice
3 Cell harvest
4 Resuspension in buffer
5 Rapid freezing
6 Cryolysis
7 Library preparation
Ribo-Seq library preparation
RNA-Seq library preparation
ATCACCGC
TAGTGGCG
8 Sequencing
CDS
9 Bioinformatic analysis

### Figure S2

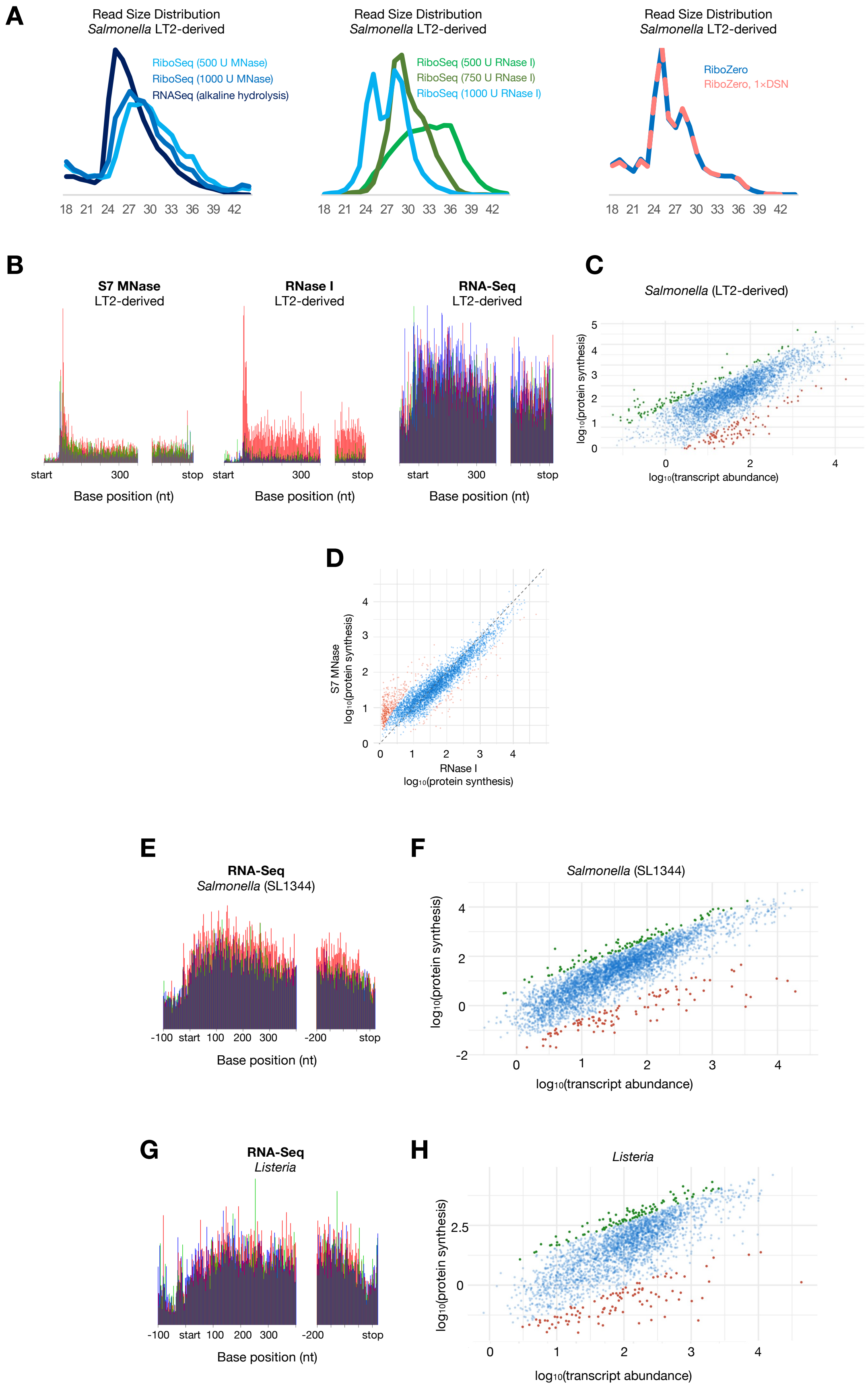
A
Read Size Distribution
Salmonella LT2-derived
RiboSeq (500 U MNase)
RiboSeq (1000 U MNase)
RNASeq (alkaline hydrolysis)
18 21 24 27 30 33 36 39 42
Read Size Distribution
Salmonella LT2-derived
RiboSeq (500 U RNase I)
RiboSeq (750 U RNase I)
RiboSeq (1000 U RNase I)
18 21 24 27 30 33 36 39 42
Read Size Distribution
Salmonella LT2-derived
RiboZero
RiboZero, 1×DSN
18 21 24 27 30 33 36 39 42
B
S7 MNase
LT2-derived
start 300 stop
Base position (nt)
RNase I
LT2-derived
start 300 stop
Base position (nt)
RNA-Seq
LT2-derived
start 300 stop
Base position (nt)
C
Salmonella (LT2-derived)
log10(protein synthesis)
log10(transcript abundance)
0 1 2 3 4 5
0 2 4
D
S7 MNase
log10(protein synthesis)
RNase I
log10(protein synthesis)
0 1 2 3 4
E
RNA-Seq
Salmonella (SL1344)
-100 start 100 200 300 -200 stop
Base position (nt)
F
Salmonella (SL1344)
log10(protein synthesis)
log10(transcript abundance)
-2 0 2 4
0 1 2 3 4
G
RNA-Seq
Listeria
-100 start 100 200 300 -200 stop
Base position (nt)
H
Listeria
log10(protein synthesis)
log10(transcript abundance)
0 2.5
0 1 2 3 4

### Figure S3

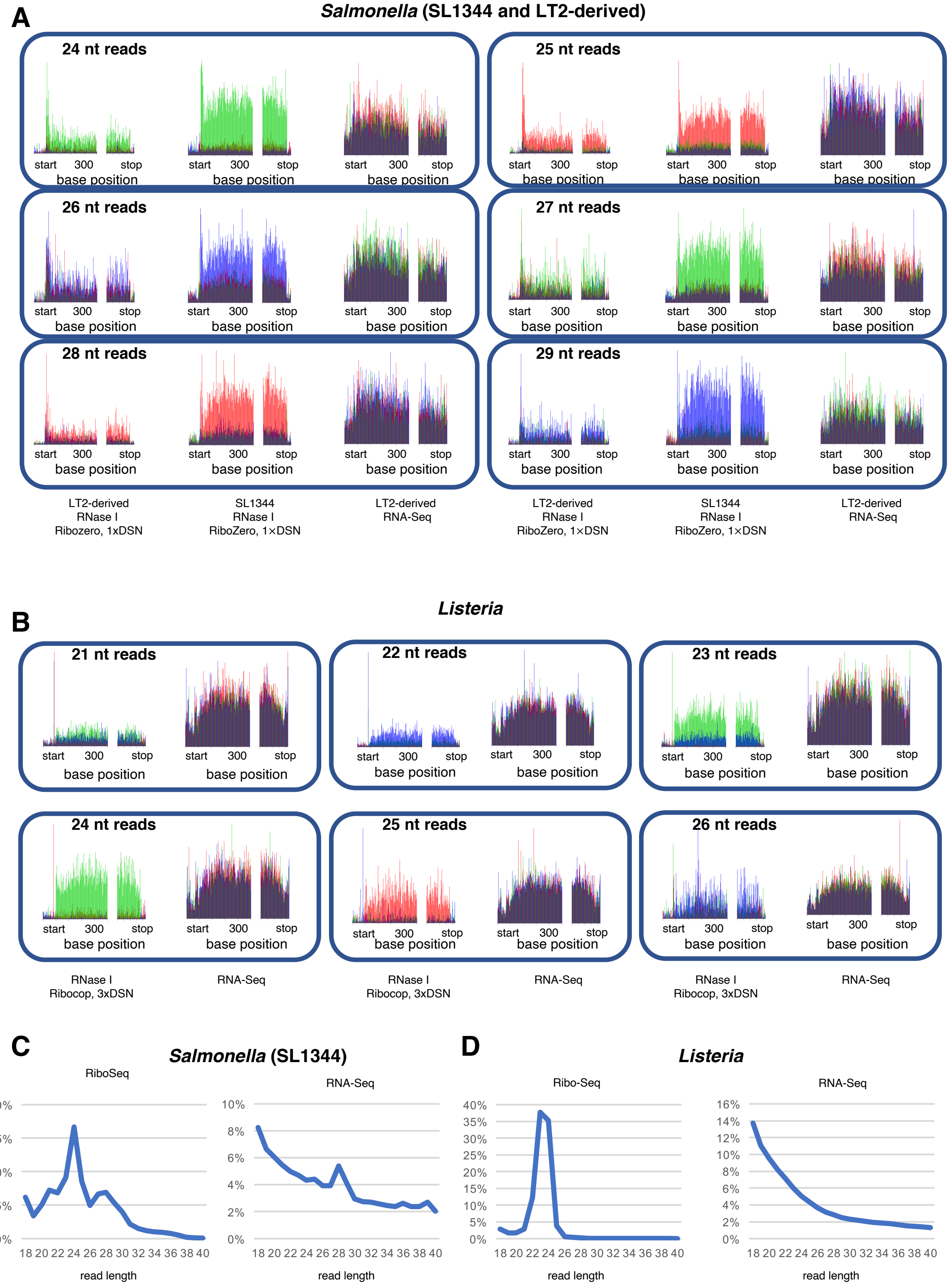
A
Salmonella (SL1344 and LT2-derived)
24 nt reads
25 nt reads
26 nt reads
27 nt reads
28 nt reads
29 nt reads
start
300
stop
base position
LT2-derived
RNase I
Ribozero, 1xDSN
SL1344
RNase I
RiboZero, 1×DSN
LT2-derived
RNA-Seq
LT2-derived
RNase I
RiboZero, 1×DSN
SL1344
RNase I
RiboZero, 1×DSN
LT2-derived
RNA-Seq
B
Listeria
21 nt reads
22 nt reads
23 nt reads
24 nt reads
25 nt reads
26 nt reads
RNase I
Ribocop, 3xDSN
RNA-Seq
C
Salmonella (SL1344)
RiboSeq
RNA-Seq
20%
15%
10%
5%
0%
8%
6%
4%
2%
18 20 22 24 26 28 30 32 34 36 38 40
read length
D
Listeria
Ribo-Seq
RNA-Seq
40%
35%
30%
25%
16%
14%
12%

### Figure S4

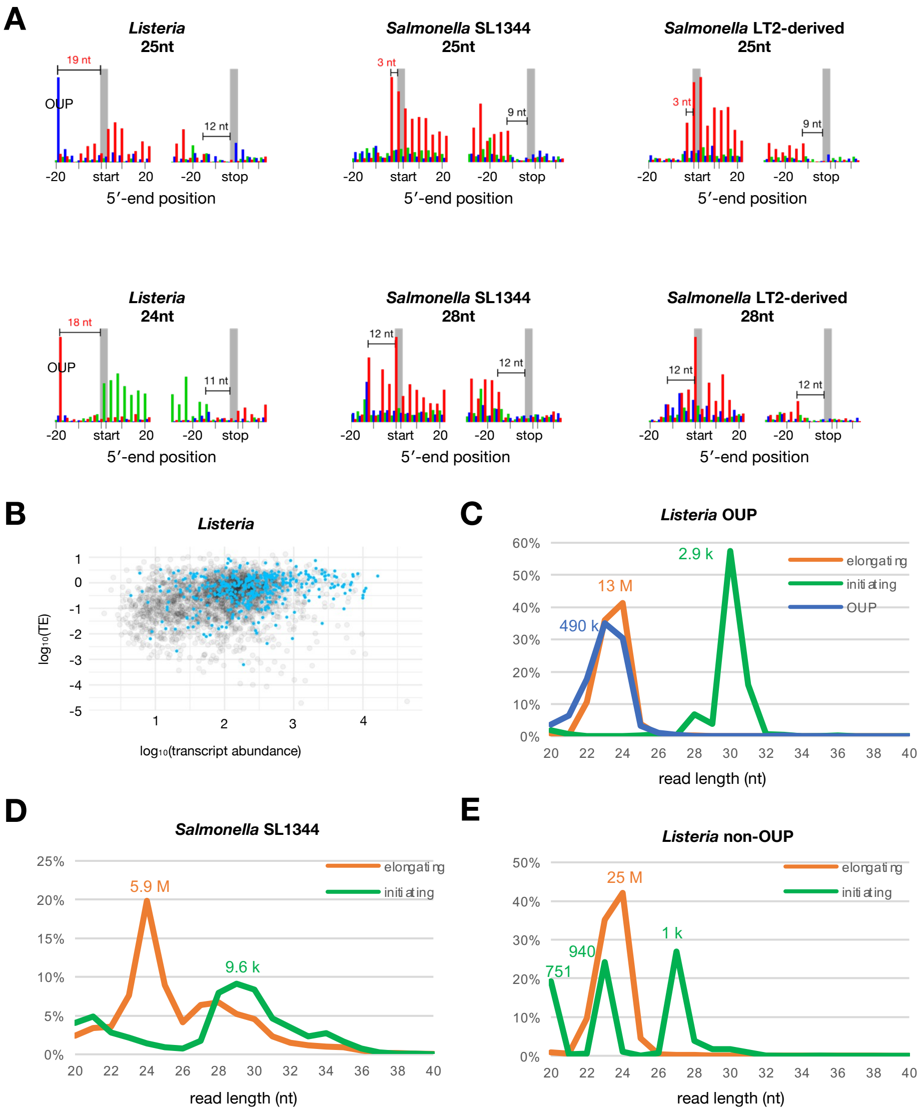
A
Listeria 25nt
19 nt
OUP
12 nt
-20 start 20 -20 stop
5′-end position
Salmonella SL1344 25nt
3 nt
9 nt
-20 start 20 -20 stop
5′-end position
Salmonella LT2-derived 25nt
3 nt
9 nt
-20 start 20 -20 stop
5′-end position
Listeria 24nt
18 nt
OUP
11 nt
-20 start 20 -20 stop
5′-end position
Salmonella SL1344 28nt
12 nt
12 nt
-20 start 20 -20 stop
5′-end position
Salmonella LT2-derived 28nt
12 nt
12 nt
-20 start 20 -20 stop
5′-end position
B
Listeria
log10(TE)
log10(transcript abundance)
C
Listeria OUP
2.9 k
13 M
490 k
elongating
initiating
OUP
read length (nt)
D
Salmonella SL1344
5.9 M
9.6 k
elongating
initiating
read length (nt)
E
Listeria non-OUP
25 M
940
751
1 k
elongating
initiating
read length (nt)

### Figure S5

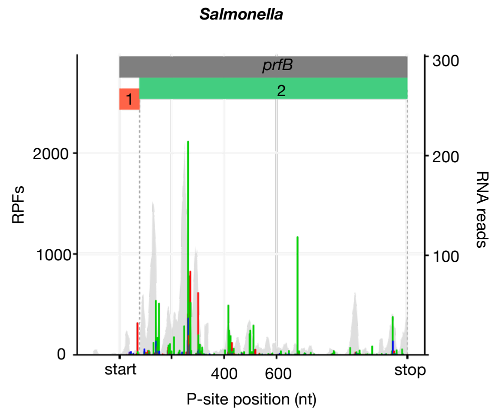

Salmonella
prfB
1
2
RPFs
RNA reads
2000
1000
0
300
200
100
start
400
600
stop
P-site position (nt)


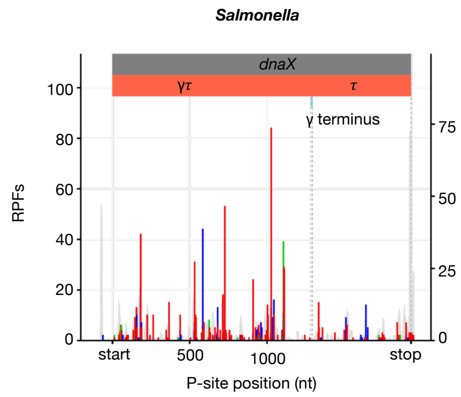

Salmonella
dnaX
γτ
τ
γ terminus
RPFs
100
80
60
40
20
0
75
50
25
0
start
500
1000
stop
P-site position (nt)


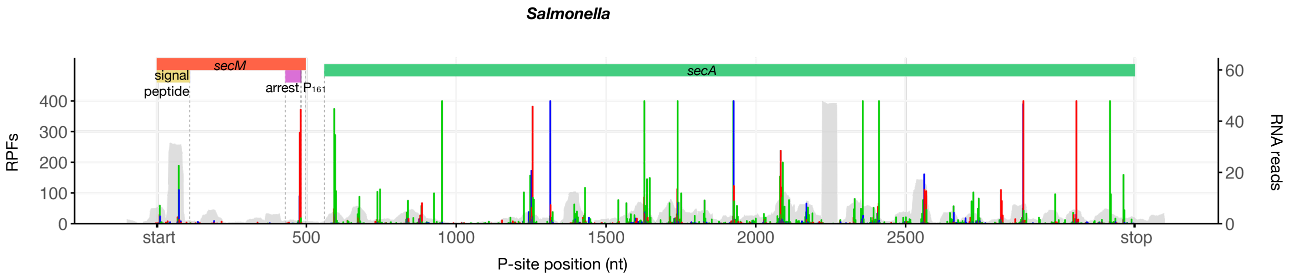

Salmonella
secM
signal
peptide
arrest
$P_{161}$
secA
RPFs
RNA reads
400
300
200
100
0
60
40
20
0
start
500
1000
1500
2000
2500
stop
P-site position (nt)

### Figure S6

**A** *Salmonella*

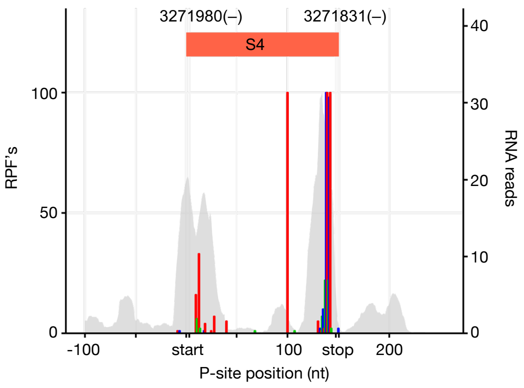


**B** *Listeria*

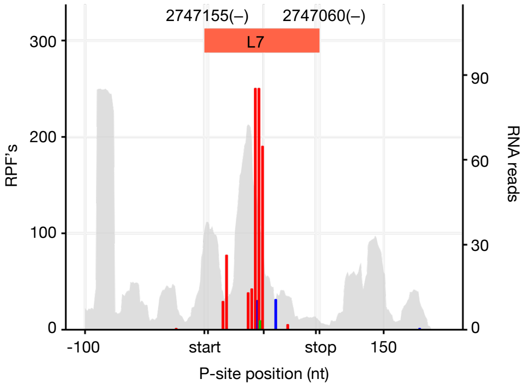

### Figure S7

**A**

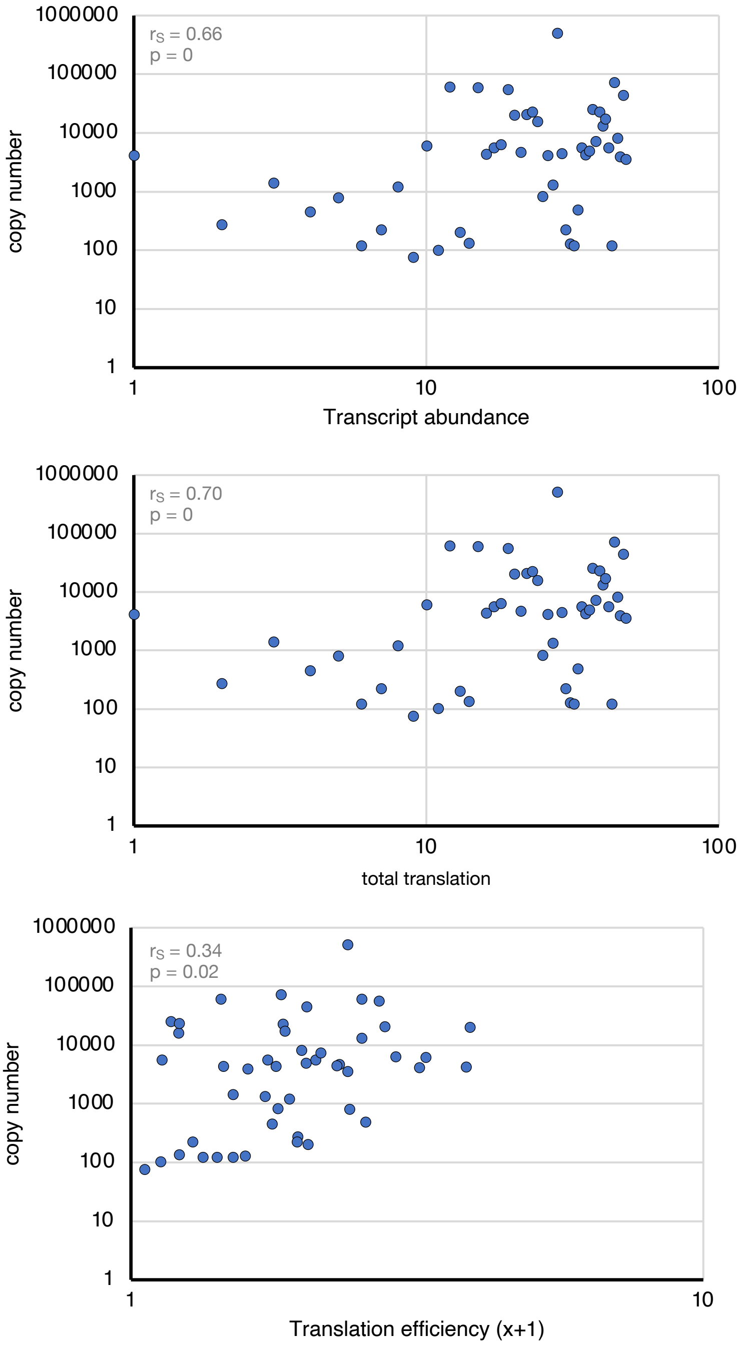
1000000
100000
10000
1000
100
10
1
copy number
$r_S$ = 0.66
p = 0
1
10
100
Transcript abundance
1000000
100000
10000
1000
100
10
1
copy number
$r_S$ = 0.70
p = 0
1
10
100
total translation
1000000
100000
10000
1000
100
10
1
copy number
$r_S$ = 0.34
p = 0.02
1
10
Translation efficiency (x+1)

### Figure S8

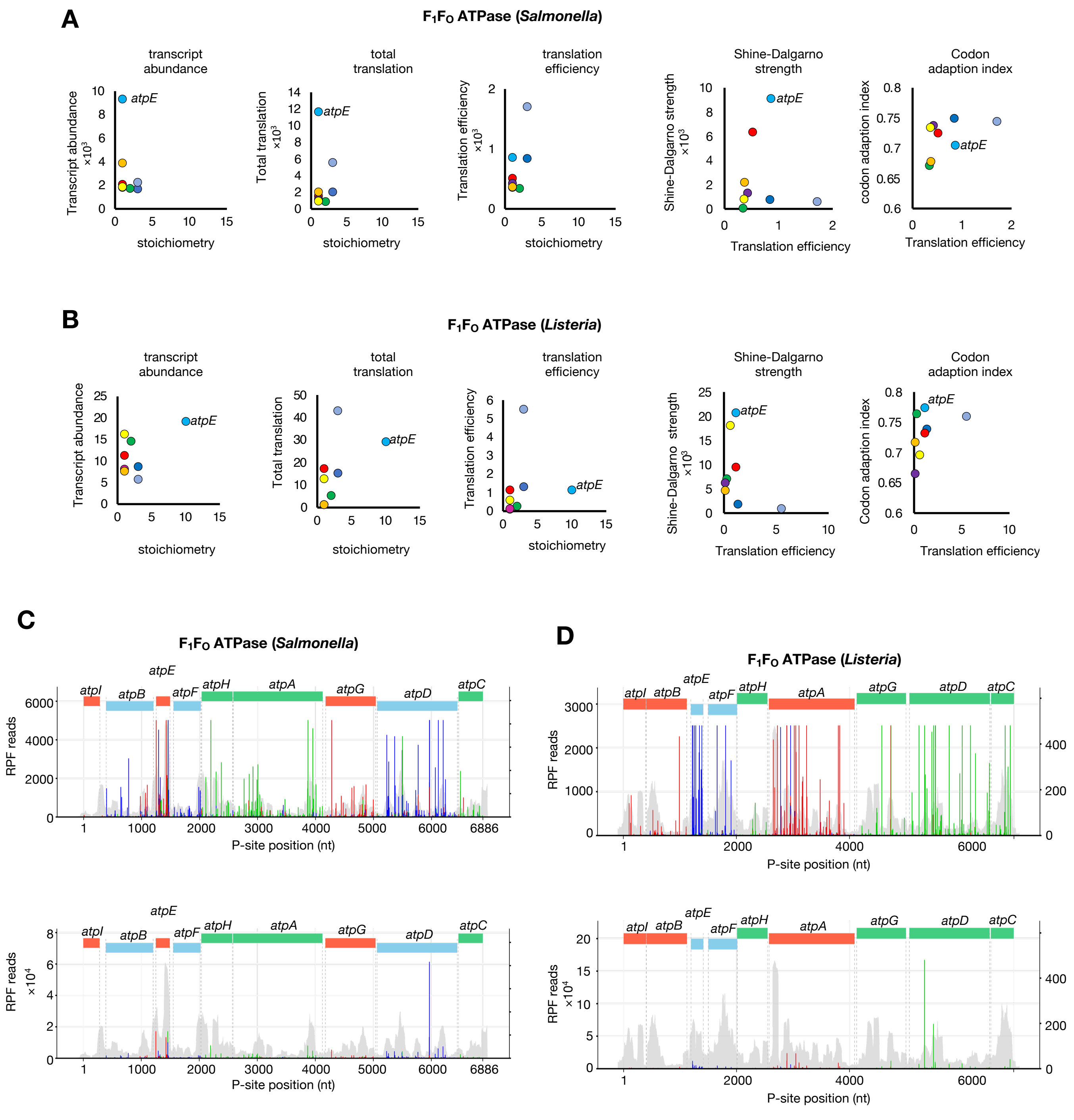

A
F1FO ATPase (Salmonella)
transcript abundance
total translation
translation efficiency
Shine-Dalgarno strength
Codon adaption index
Transcript abundance ×10^3
Total translation ×10^3
Translation efficiency ×10^3
Shine-Dalgarno strength ×10^3
codon adaption index
stoichiometry
Translation efficiency
atpE
B
F1FO ATPase (Listeria)
Transcript abundance
Total translation
Translation efficiency
Shine-Dalgarno strength ×10^3
Codon adaption index
C
F1FO ATPase (Salmonella)
D
F1FO ATPase (Listeria)
atpI
atpB
atpE
atpF
atpH
atpA
atpG
atpD
atpC
RPF reads
RPF reads ×10^4
P-site position (nt)

### Figure S9

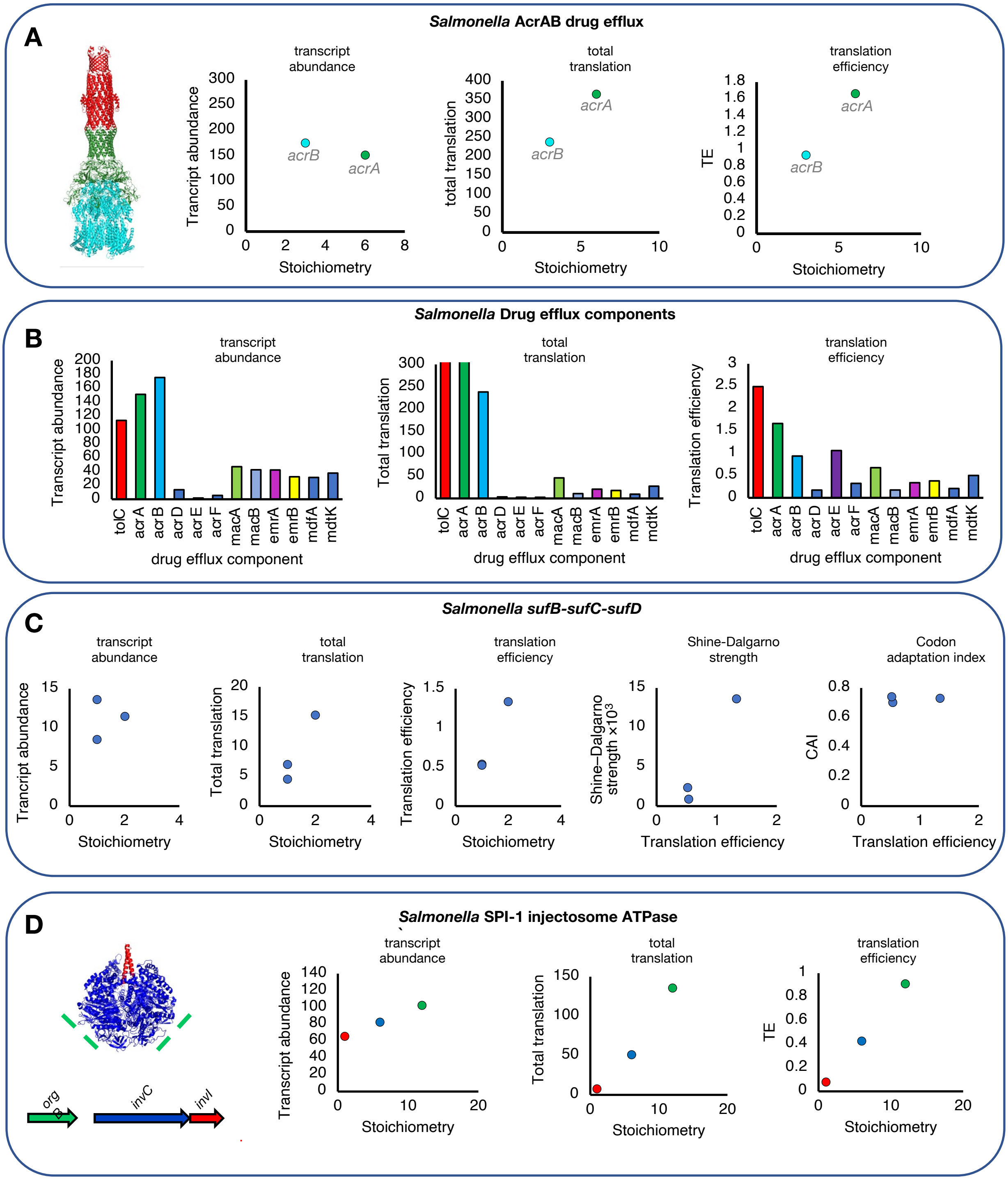
A
*Salmonella* AcrAB drug efflux
transcript abundance
Trancript abundance
Stoichiometry
acrB
acrA
total translation
total translation
translation efficiency
TE
B
*Salmonella* Drug efflux components
Transcript abundance
Total translation
Translation efficiency
tolC
acrA
acrB
acrD
acrE
acrF
macA
macB
emrA
emrB
mdfA
mdtK
drug efflux component
C
*Salmonella sufB-sufC-sufD*
Shine-Dalgarno strength
Shine–Dalgarno strength ×10³
Codon adaptation index
CAI
Translation efficiency
D
*Salmonella* SPI-1 injectosome ATPase
orgB
invC
invI

### Figure S10
### LISTERIA

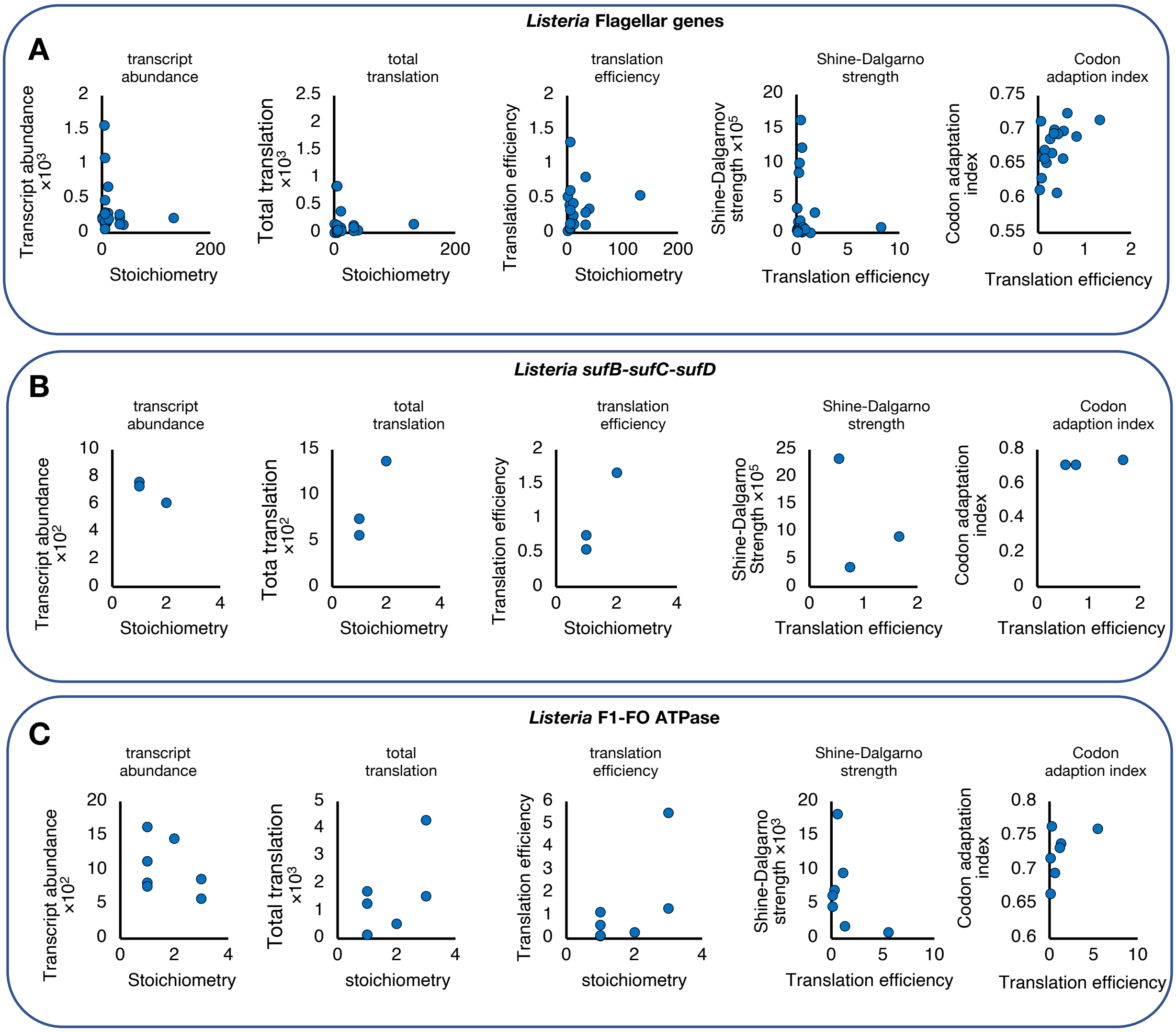

### Figure S11

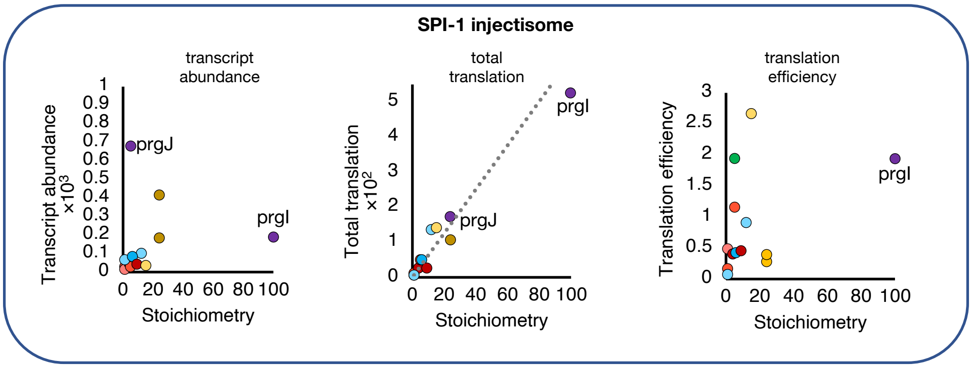
SPI-1 injectisome
transcript abundance
Transcript abundance ×10³
1
0.9
0.8
0.7
0.6
0.5
0.4
0.3
0.2
0.1
0
prgJ
prgI
0 20 40 60 80 100
Stoichiometry
total translation
Total translation ×10²
5
4
3
2
1
0
prgI
prgJ
0 20 40 60 80 100
Stoichiometry
translation efficiency
Translation efficiency
3
2.5
2
1.5
1
0.5
0
prgI
0 20 40 60 80 100
Stoichiometry

### Figure S12

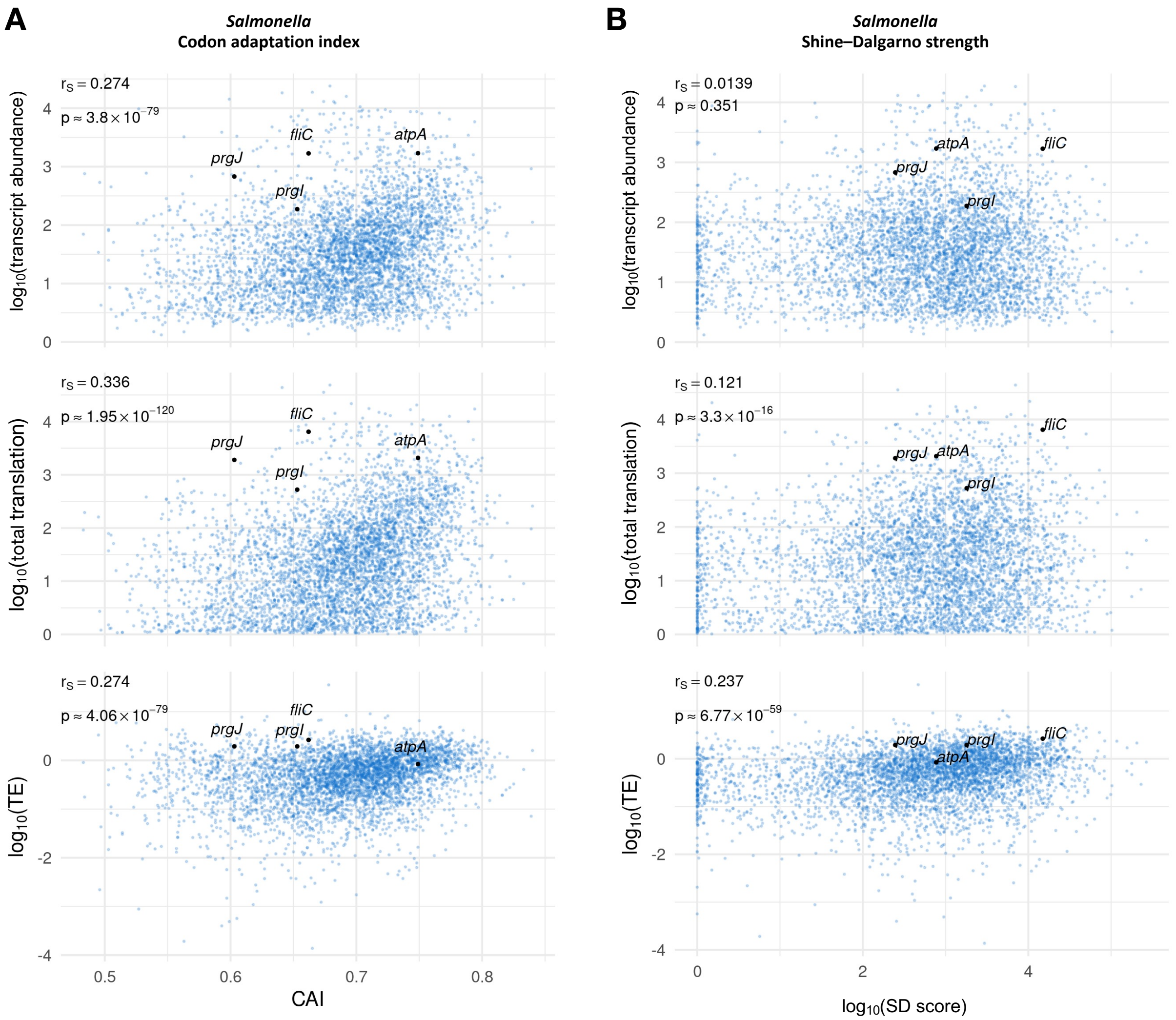
A
Salmonella
Codon adaptation index
B
Salmonella
Shine–Dalgarno strength
rS = 0.274
p ≈ 3.8 × 10^−79
rS = 0.336
p ≈ 1.95 × 10^−120
rS = 0.274
p ≈ 4.06 × 10^−79
rS = 0.0139
p ≈ 0.351
rS = 0.121
p ≈ 3.3 × 10^−16
rS = 0.237
p ≈ 6.77 × 10^−59
fliC
atpA
prgJ
prgI
log10(transcript abundance)
log10(total translation)
log10(TE)
CAI
log10(SD score)

### Figure S13

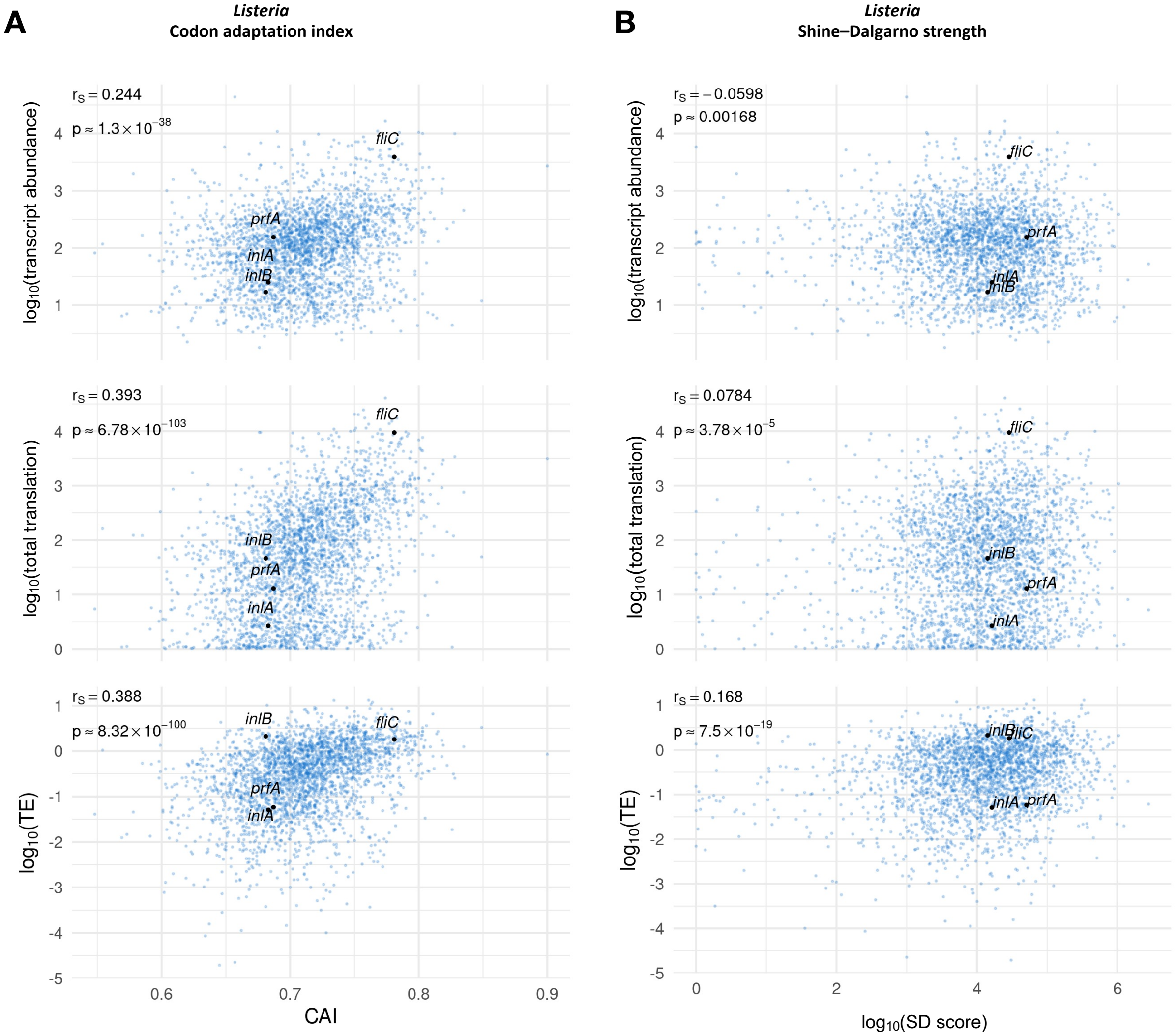
A
Listeria
Codon adaptation index
B
Listeria
Shine–Dalgarno strength
$r_S = 0.244$
$p \approx 1.3 \times 10^{-38}$
$r_S = -0.0598$
$p \approx 0.00168$
$r_S = 0.393$
$p \approx 6.78 \times 10^{-103}$
$r_S = 0.0784$
$p \approx 3.78 \times 10^{-5}$
$r_S = 0.388$
$p \approx 8.32 \times 10^{-100}$
$r_S = 0.168$
$p \approx 7.5 \times 10^{-19}$
$\log_{10}$(transcript abundance)
$\log_{10}$(total translation)
$\log_{10}$(TE)
fliC
prfA
inlA
inlB
CAI
$\log_{10}$(SD score)

### Figure S14 NEW

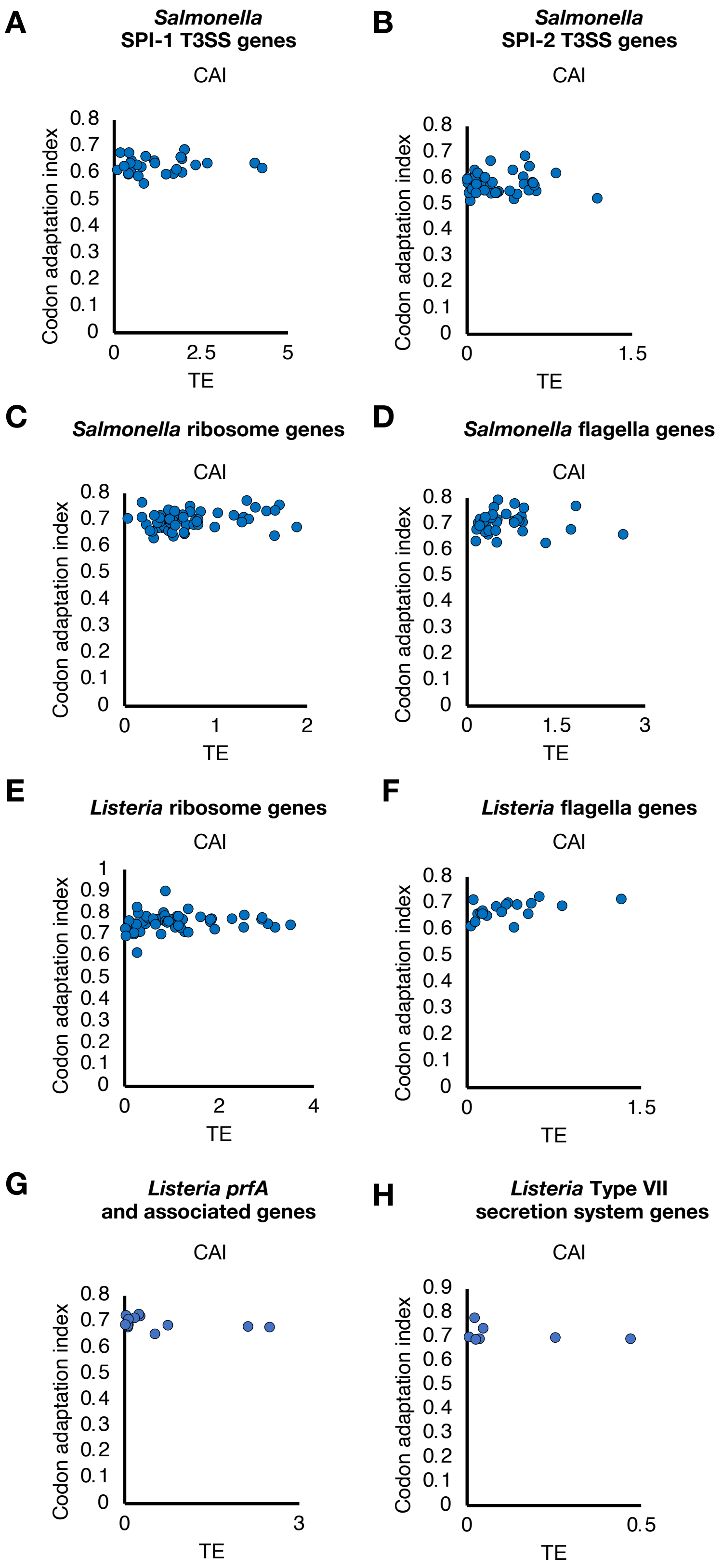

### Figure S15 NEW

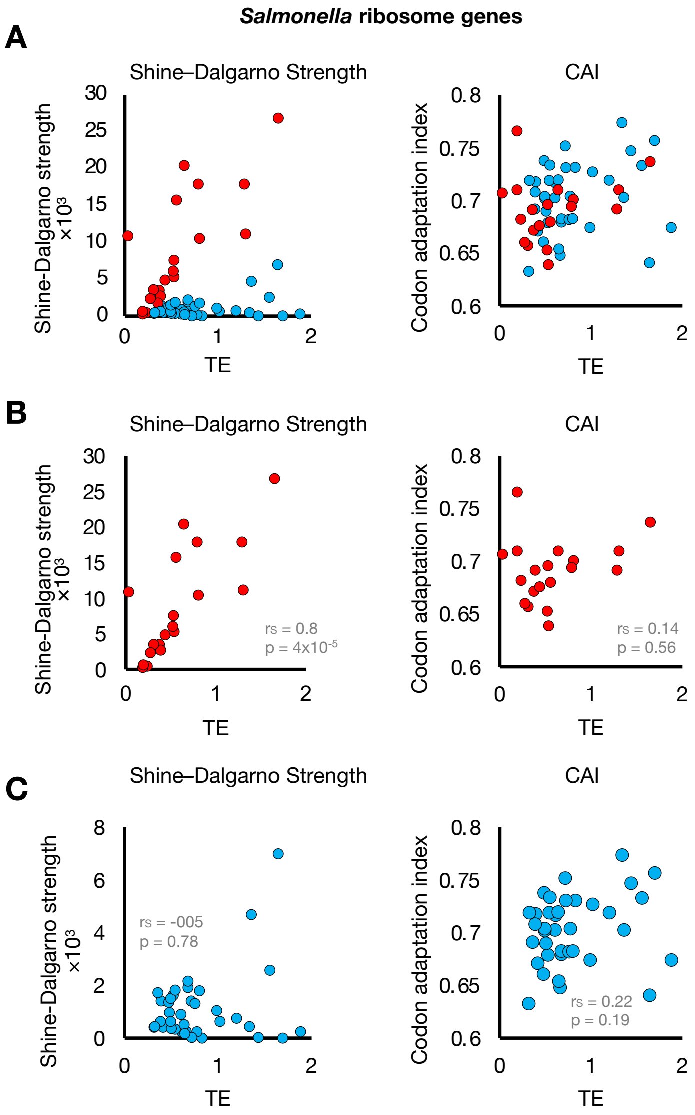

### Figure S16 NEW

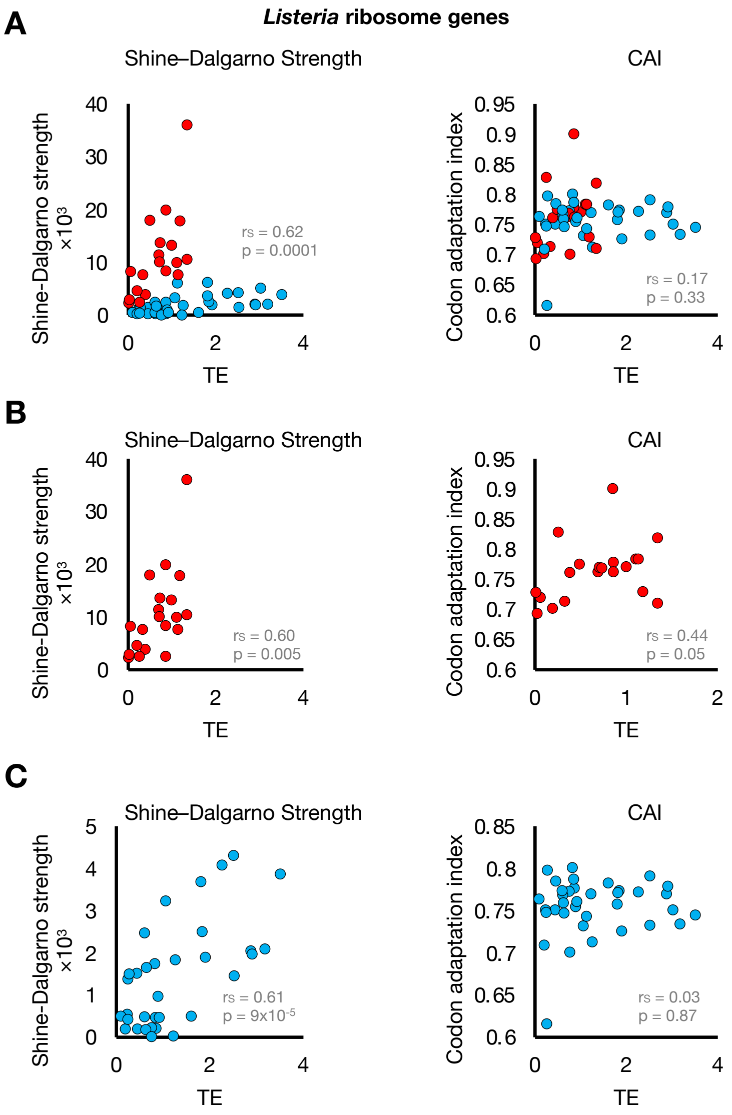

### Figure S17 NEW

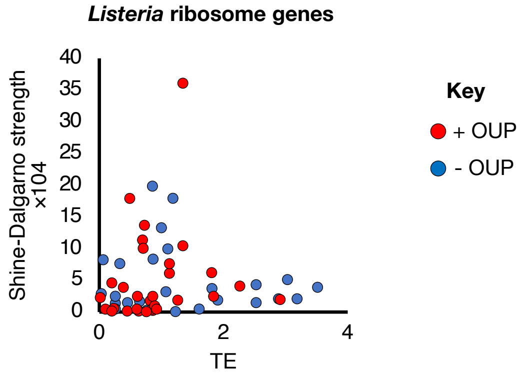

### Figure S18

***Salmonella***

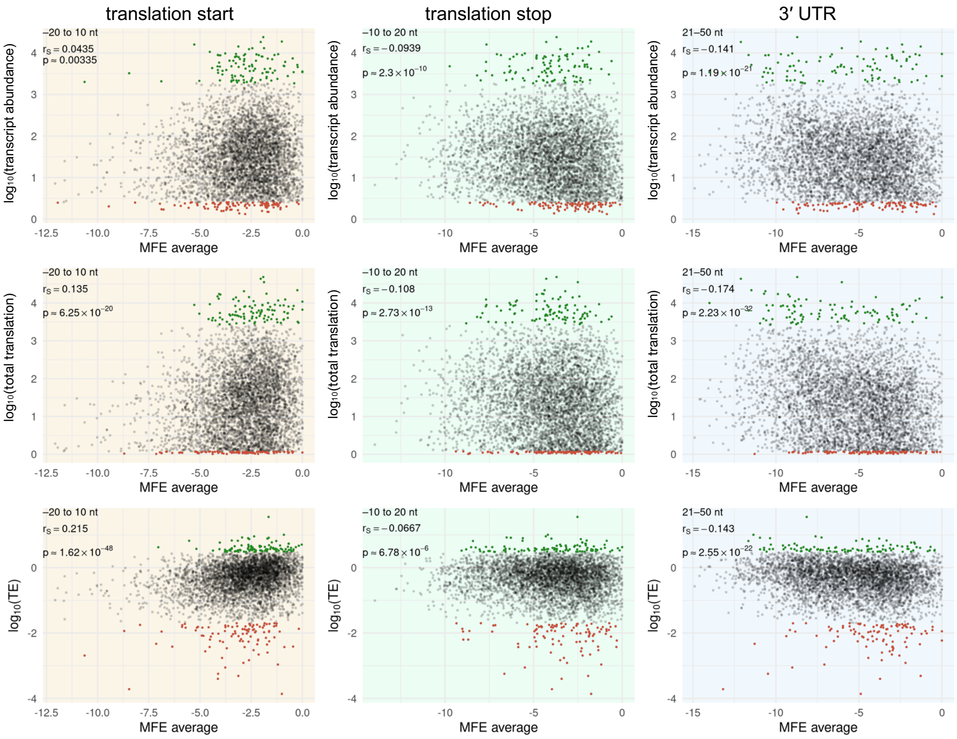


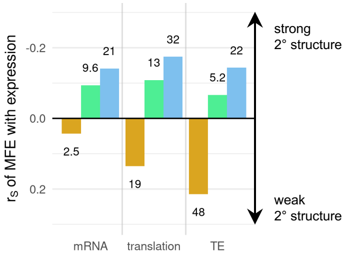

### Figure S19

***Listeria***

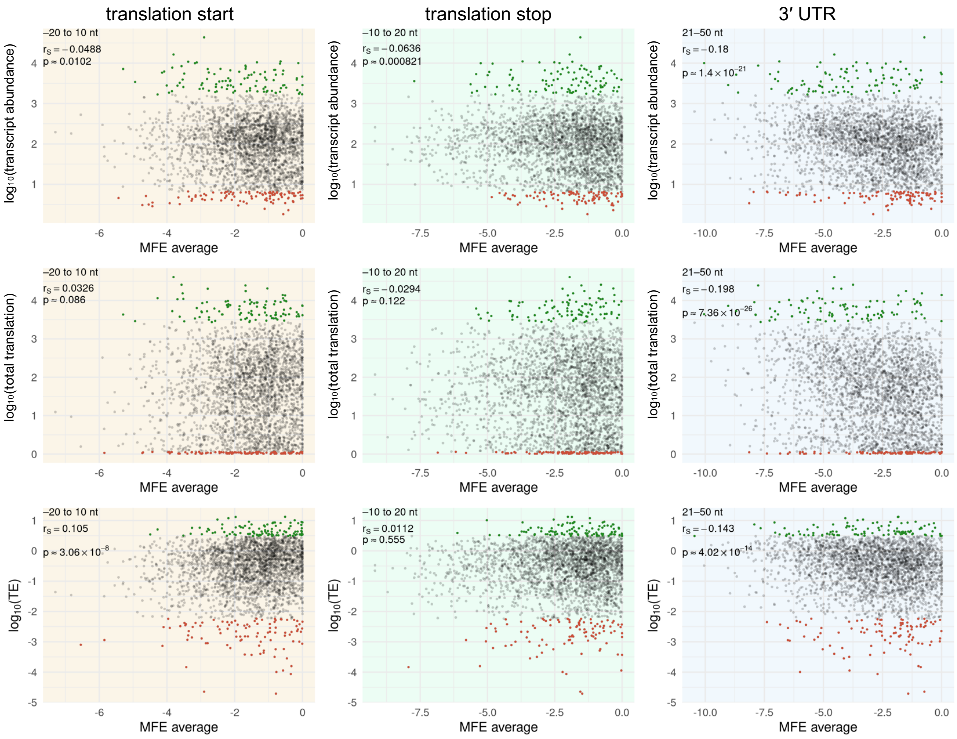


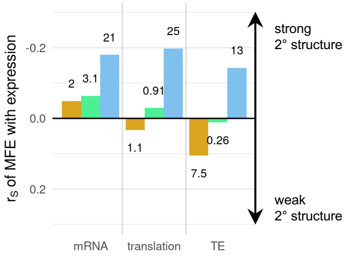

### Figure S20

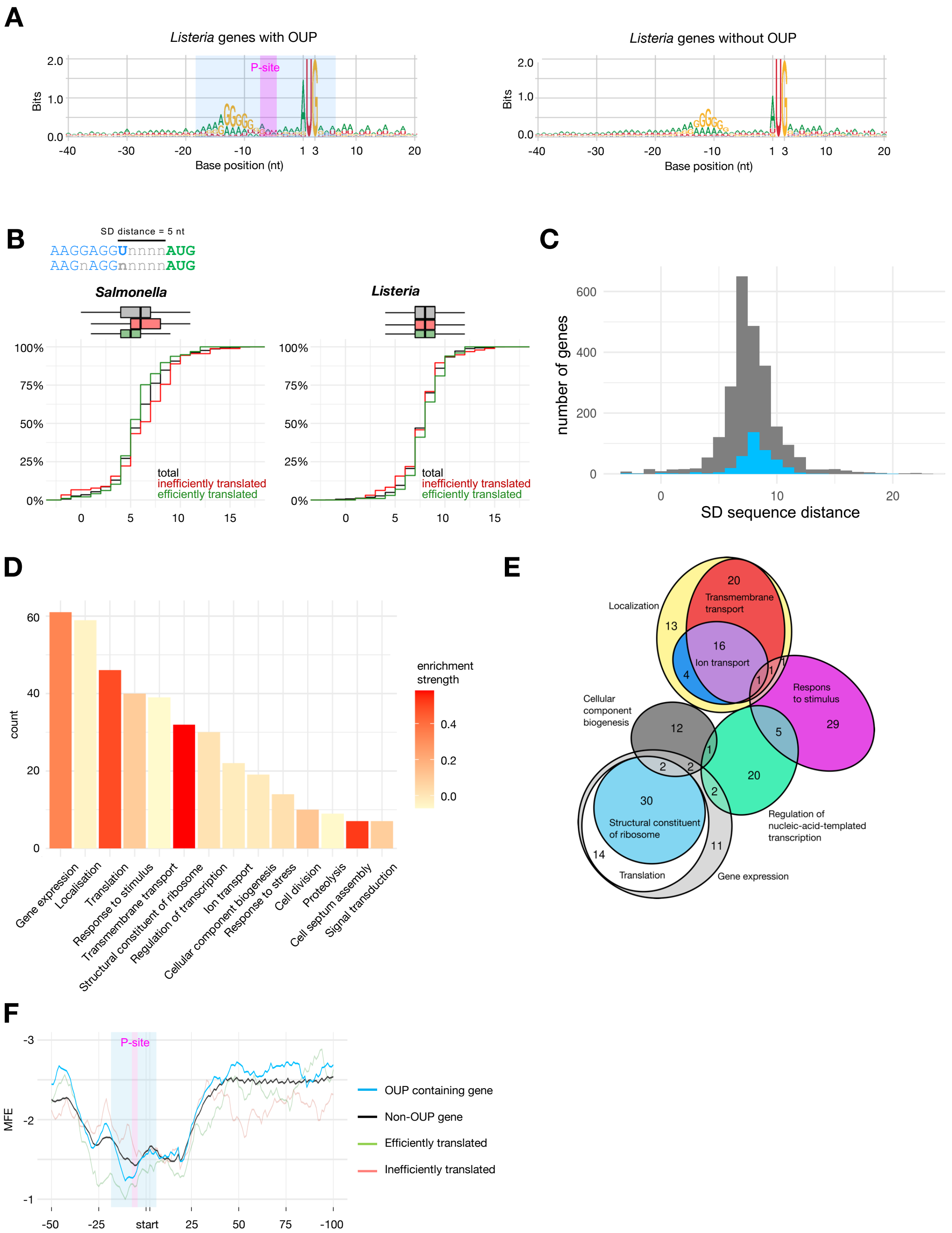
A
Listeria genes with OUP
Listeria genes without OUP
P-site
Bits
Base position (nt)
B
SD distance = 5 nt
AAGGAGGUnnnnAUG
AAGnAGGnnnnnAUG
Salmonella
Listeria
total
inefficiently translated
efficiently translated
C
number of genes
SD sequence distance
D
count
enrichment strength
Gene expression
Localisation
Translation
Response to stimulus
Transmembrane transport
Structural constituent of ribosome
Regulation of transcription
Ion transport
Cellular component biogenesis
Response to stress
Cell division
Proteolysis
Cell septum assembly
Signal transduction
E
Localization
Transmembrane transport
Ion transport
Respons to stimulus
Cellular component biogenesis
Structural constituent of ribosome
Regulation of nucleic-acid-templated transcription
Translation
Gene expression
F
P-site
MFE
start
OUP containing gene
Non-OUP gene
Efficiently translated
Inefficiently translated

### Figure S21

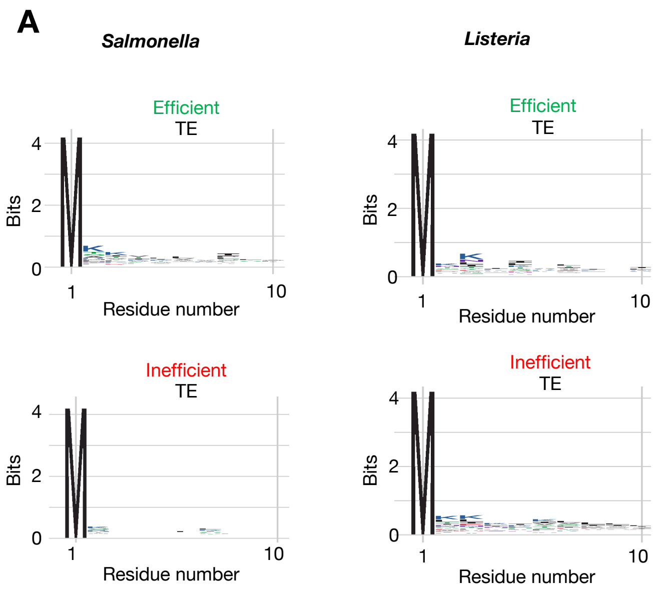

A
Salmonella
Listeria
Efficient
TE
Inefficient
TE
Bits
Residue number
4
2
0
1
10


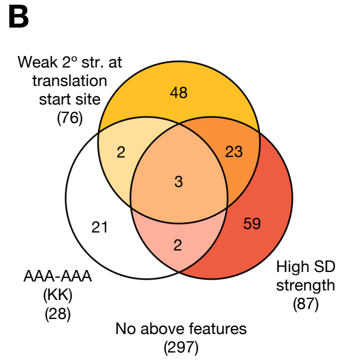

B
Weak 2º str. at translation start site (76)
48
2
23
3
21
59
2
AAA-AAA (KK) (28)
High SD strength (87)
No above features (297)


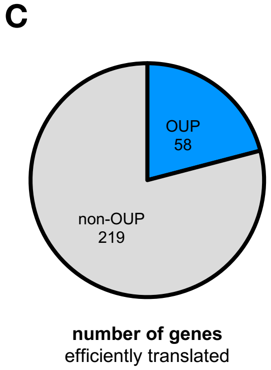

C
OUP
58
non-OUP
219
number of genes
efficiently translated


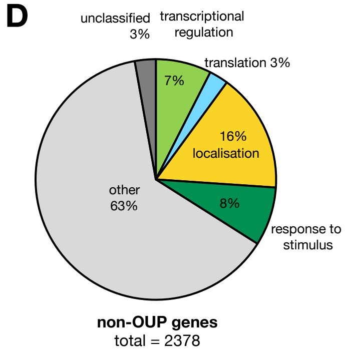

D
unclassified 3%
transcriptional regulation
7%
translation 3%
16% localisation
8%
response to stimulus
other 63%
non-OUP genes
total = 2378

### Figure S22

**A**

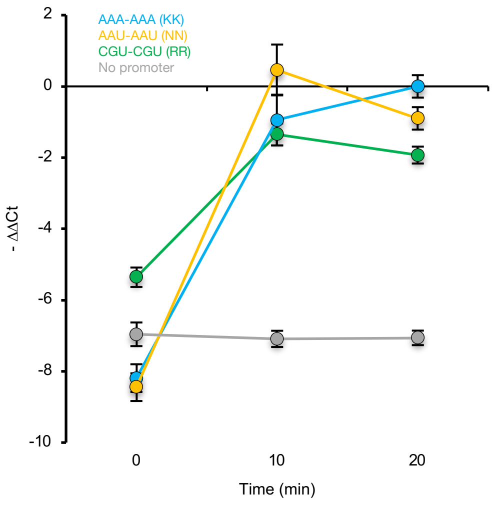
AAA-AAA (KK)
AAU-AAU (NN)
CGU-CGU (RR)
No promoter
2
0
-2
-4
-6
-8
-10
- ΔΔCt
0
10
20
Time (min)
